# VULCAN integrates ChIP-seq with patient-derived co-expression networks to identify GRHL2 as a key co-regulator of ERa at enhancers in breast cancer

**DOI:** 10.1101/266908

**Authors:** Andrew N. Holding, Federico M. Giorgi, Amanda Donnelly, Amy E. Cullen, Sankari Nagarajan, Luke A Selth, Florian Markowetz

**Affiliations:** CRUK Cambridge Institute, University of Cambridge, Robinson Way, Cambridge, CB2 0RE; The Alan Turing Institute, 96 Euston Road, Kings Cross, London NW1 2DB; Department of Pharmacy and Biotechnology, University of Bologna, Via Selmi 3, Bologna, Italy; Dame Roma Mitchell Cancer Research Laboratories and Freemasons Foundation Centre for Men’s Health, Adelaide Medical School, The University of Adelaide, SA, Australia

**Keywords:** Breast Cancer, Network Analysis, Dynamics, ER, Master Regulator, ChIP-seq, VULCAN, GRHL2, P300, H3K27ac

## Abstract

**Background:** **V**irt**U**a**L C**hIP-seq **A**nalysis through **N**etworks (VULCAN) infers regulatory interactions of transcription factors by overlaying networks generated from publicly available tumor expression data onto ChIP-seq data. We applied our method to dissect the regulation of Estrogen Receptor-alpha (ER) activation in breast cancer to identify potential coregulators of the ER’s transcriptional response.

**Results:** VULCAN analysis of ER activation in breast cancer highlighted key components of the ER complex alongside a novel interaction with GRHL2. We demonstrate that GRHL2 is recruited to a subset of ER binding sites and regulates the transcriptional output of ER, as evidenced by changes in ER-associated eRNA expression, and stronger ER binding at active enhancers (H3K27ac sites) after GRHL2 knockdown.

**Conclusions:** Our findings provide new insight into the role of GRHL2 in regulating eRNA transcription as part of ER signaling. These results demonstrate VULCAN, available from Bioconductor, as a powerful predictive tool.

## Introduction

Breast cancer is the most common form of cancer in women in North America and Europe accounting for 31% of all new cancer cases. In the US, it is estimated that 41,400 deaths will have occurred from the disease in 2018 [1]. The majority of breast cancers, approximately 70%, are associated with deregulated signaling by the Estrogen Receptor-alpha (ER), which drives tumor growth. Therefore, in ER-positive (ER+) tumors, ER is the primary therapeutic target. During activation, ER recruits several cofactors to form an active complex on the chromatin. FOXA1 is of particular interest as the protein shares nearly 50% of its genomic binding sites with ER and has been shown to operate as a pioneer factor before ER activation [2], [3]. It is through FOXA1 and other cofactors (e.g SRC-1) [4], [5], that ER is able to recruit RNA Polymerase II at the gene promoter sites by way of adaptor proteins in order to initiate transcription [6]. Combinatorial treatments targeting ER cofactors present a significant opportunity in breast cancer therapy for increasing patient survival. In particular, the pioneer factor FOXA1 [7] has been identified as novel therapeutic target for the treatment of breast cancer, while the EZH2-ERα-GREB1 transcriptional axis has been shown to play a key role in therapeutic resistance [8].

ChIP-seq enables the identification of potential site-specific interactions at common binding sites between transcription factors and their cofactors; however, to fully characterize all potential cofactors of a single project on this scale is laborious and expensive. To follow up all potential cofactors identified by a chromatin-wide proteomics method, e.g. RIME [9] or ChIP-MS [10], would take hundreds of individual ChIP-seq experiments. Studies like ENCODE [11] have gone a long way to provide a resources to meet these challenges; however, the inherent scale of the problem means public studies can only offer data for a subset of TF in a limited number of models. For a single lab to undertake this level of experimentation is unfeasible, and in cases where suitable antibodies for the ChIP do not exist, impossible.

To enable discoveries beyond collections like ENCODE, we are proposing a computational framework to integrate patient data in the the prediction of functional protein-protein interactions. By applying machine learning methods, we are able to surpass the limitation of current predictive tools that exist to support the interpretation of data. Previous methods provide information in the context of predefined biological pathways and established gene sets [12], [13] or through motif analysis [14], while our method is built on data specific to the disease being studied. Further, standard gene set enrichment analysis has inherent limitations because it was not designed for reconstructing gene networks, whereas one of the advantages of VULCAN is that it downweights genes shared by multiple TFs.

Our method, “**V**irt**U**a**L C**hIP-seq **A**nalysis through **N**etworks” (VULCAN), is able to specifically analyze potential disease-specific interactions of TFs in ChIP-seq experiments by combining machine learning approaches and patient data. Previously, the strategies employed by VULCAN were limited to the analysis of transcription data. By developing VULCAN to overlay coexpression networks established from patient tumor data onto ChIP-seq data, we are able to provide candidate coregulators of the response to a given stimulus (Fig. 1). Further, as VULCAN builds on transcriptional master regulator analysis, the output from the pipeline provides the end user with functional information in terms of the activity of potentially interacting TFs. The combination of disease-specific context and TF activity information presents a significant step forward in providing valuable information for the elucidation of on-chromatin interactions from ChIP-seq experiments over previous strategies.

**Figure 1:**
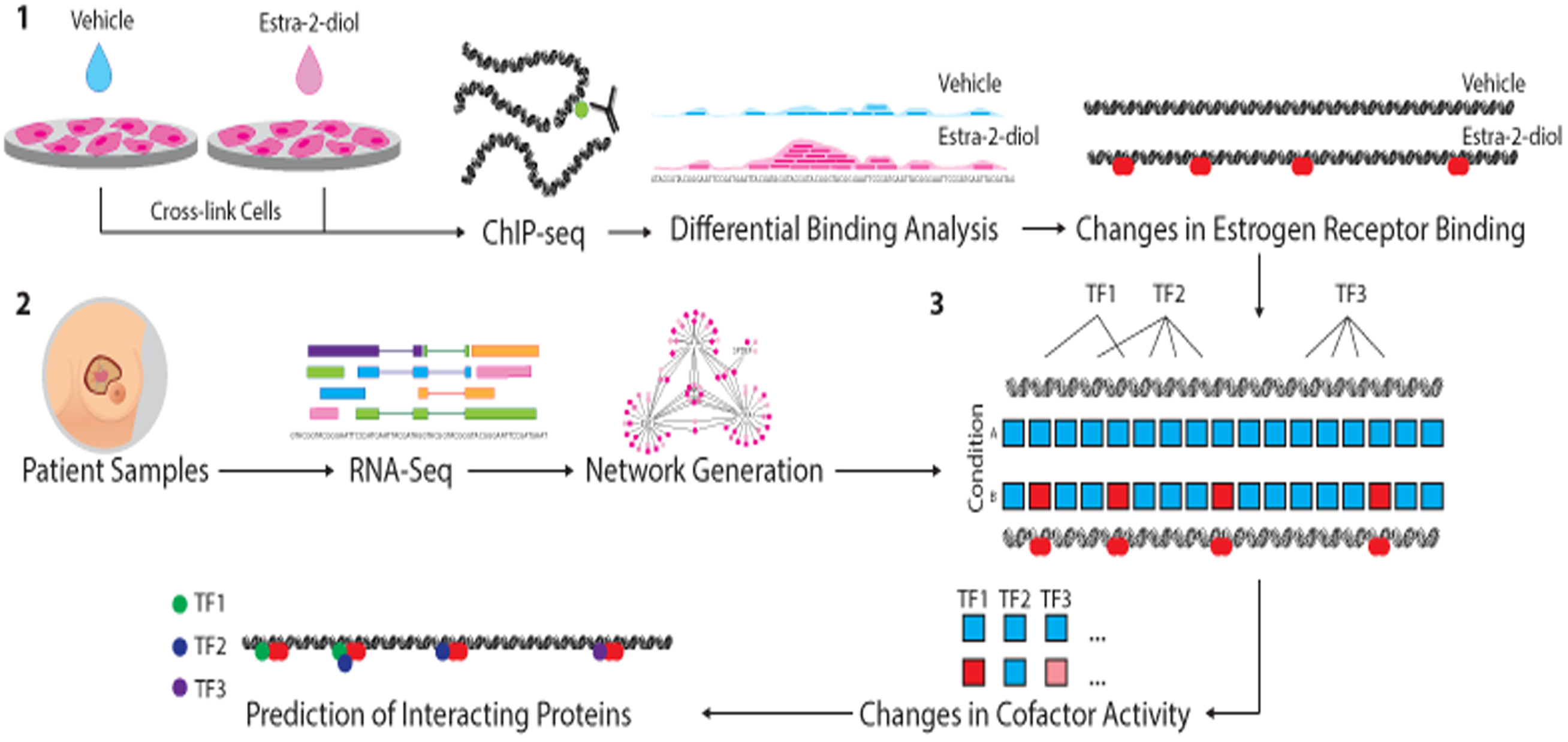
An overview of VULCAN. **1.** ChIP-seq analysis from multiple conditions is undertaken to generate cistrome data at multiple timepoints (or conditions). Binding events are then compared using differential binding analysis to establish log-fold change values for individual binding events between each timepoint. **2.** Network generation was undertaken with ARACNe-AP by inferring all pairwise TF-target coexpression from patient datasets (e.g. TCGA breast & METABRIC datasets). **3.** All the targets of each specific TF in the network, i.e. the individual regulons, are tested against the established changes in ER binding through the msVIPER algorithm [15] to identify proteins that interact with the target transcriptional factor and final prediction is given for potential interacting cofactors.

Through the application of VULCAN to the activation of the ER in breast cancer, we were able to identify multiple previously characterized cofactors of the ER along with GRHL2 as a potential co-repressor of the ER. We then demonstrated experimentally that GRHL2 is able to modulate the expression of eRNA at ER bound enhancers, and the removal of the P300 inhibitory alpha-helix results in suppression of the inhibitory effect on eRNA production.

## Results

VULCAN integrates ChIP-seq data (Fig. 1**, Step 1**) with co-expression networks (Fig. 1**, Step 2**) to predict cofactor activity (Fig. 1**, Step 3**). The initial ChIP-seq data is converted into genomic regions, and if multiple conditions are supplied the changes in the transcription factor affinity are calculated. In parallel, master regulator analysis of tumour transcriptional data is used to provide tissue specific information on the regulation of genes by TFs within the tumour type. The integration of these two data types provides context-specific results and differentiates VULCAN from the existing methods which make use of predefined unweighted gene sets or motif analysis. VULCAN additionally makes use of key the functionality of the VIPER algorithm [15] that assigns edge-specific scores like Mode of Action and likelihood to the reconstructed network.

In the following, we first benchmark VULCAN’s performance in a comprehensive comparison to alternative approaches. We then apply it to our data on temporal ER binding, which identifies GRHL2 as a novel ER cofactor and we explore its function.

### Comparison of VULCAN to existing methods

#### Mutual Information networks outperform Partial Correlation networks

We generated a mutual information network with ARCANe alongside several partial correlation networks at different thresholds all from the TCGA Breast Cancer data. To ensure robustness of our method, we tested the overlap of every partial correlation network with the mutual information network using the Jaccard Index (JI) criterion (**Suppl. Fig. S1**). Finally, we showed how the Jaccard Index between partial correlation networks and the ARACNe network is always significantly higher than expected by selecting random network edges (**Suppl. Fig. S2**). For further analysis, we selected the mutual information network generated by ARACNe as this method outperformed partial correlation networks at all thresholds.

#### GSEA is the optimum method for VULCAN’s target enrichment analysis

VULCAN applies Gene Set Enrichment Analysis [16] to identify enrichment of our mutual information network derived regulons in differential ChIP-seq data. To validate our method, we compared the results of VULCAN when applied to our ER binding data against three independent methods previously applied to benchmark VIPER [15]. The first implemented a Fisher p-value integration step. This test lacks stringency and results in nearly all regulons as significantly enriched (**Suppl. Fig. S3**). Second, we implemented a fraction of targets method, defining for every TF the fraction of their targets that are also differentially bound. This alternative to VULCAN ignores the MI strength of interaction and the individual strengths of differential bindings, reducing the resolving power of the algorithm (**Suppl. Fig. S4**). Finally, we compared to a Fisher’s Exact Method, which assesses the overlap between networks and significant differential binding. This method is too stringent (as observed in the original VIPER paper) [15]; and even without p-value correction, there are no significant results, even at low stringency, demonstrating the low sensitivity of using a Fisher’s Exact Method method (**Suppl. Fig. S5**). In summary, VULCAN GSEA implementation outperformed all three alternative methods we tested (t-test based; fraction of targets method; and Fisher’s Exact Method) in our dataset and was therefore applied to all downstream analysis of ChIP-seq data.

#### VULCAN outperforms enrichment analysis tools (GREAT, ISMARA & ChIP-Enrich)

To further validate our method, we compared the output of our GSEA analysis with different versions of promoter-enrichment approaches implemented by GREAT [12], ISMARA [14] and ChIP-Enrich [13]. The VULCAN analysis shows a significant overlap in terms of detected pathways with the GREAT method (**Suppl. Fig. S6**). ChIP-Enrich identifies enrichment of a number of TFs also predicted by VULCAN, but it fails to identify ESR1 as the top transcription factor affected by our experiment (**Suppl. Fig. S7**). ISMARA succeeds at identifying ESR1 using a motif-based analysis, but does not identify other candidate binding TFs (**Suppl. Fig. S8**). In summary, VULCAN outperforms both ISMARA and ChIP-Enrich, and significantly overlaps with GREAT, but provides additional value through inference of TF factor activity.

### Temporal analysis of ER DNA binding profiles after activation by E2

We performed four replicated ChIP-seq experiments for ER at three time points (0, 45 and 90 minutes) after estradiol treatment (Fig. 2) in the ER+ breast cancer cell line, MCF7. The cistromic profile of ER at 45 and 90 minutes was then compared to 0 minutes to identify binding events enriched by E2. Our analysis (Fig. 2B,C) identified 18,900 statistically significant binding events at 45 minutes (FDR < 0.05) and 17,896 numbers at 90 minutes. We validated the ER binding behavior with ChIP-qPCR (Fig. 2A) and the response was sustained, in agreement with our previous study. [17]

**Figure 2:**
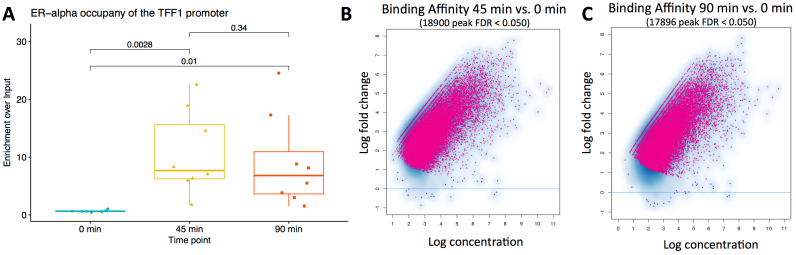
Dynamic behavior during early activation of ER. ChIP-qPCR of the TFF1 gene (A) at 3 timepoints shows increased binding of ER at 45 minutes after MCF7 cells are stimulated by estradiol. The previously reported maximum is followed by a decrease in the TFF1 promoter occupancy at 90 minutes. P-values are generated by one-tailed t-test. The maximal point at 90 minutes was identified as an outlier (> median + 2 × IQR); however, removal did not alter the significance of results. (B) Differential binding analysis of ChIP-seq data at three timepoints to monitor the activation of ER. The ER exhibits a strong increase in binding at 45 minutes vs 0 minutes (C) and the majority of sites still display binding at 90 minutes.

We performed motif enrichment analysis (HOMER software) on ER binding sites detected by differential binding analysis. Our analysis confirmed a strong enrichment for a single element, ERE, bound at both 45 and 90 minutes, with a corrected p-value of 0.0029 (Fig. 3F). When clustered according to peak intensity, samples cluster tightly in two groups: treated and untreated (**Suppl. Fig. S9, S10 and S11**), but treatment at 45 and 90 minutes is detectably different on a genome-wide scale, as highlighted by Principal Component Analysis (**Suppl. Fig. S12 and S13**).

**Figure 3:**
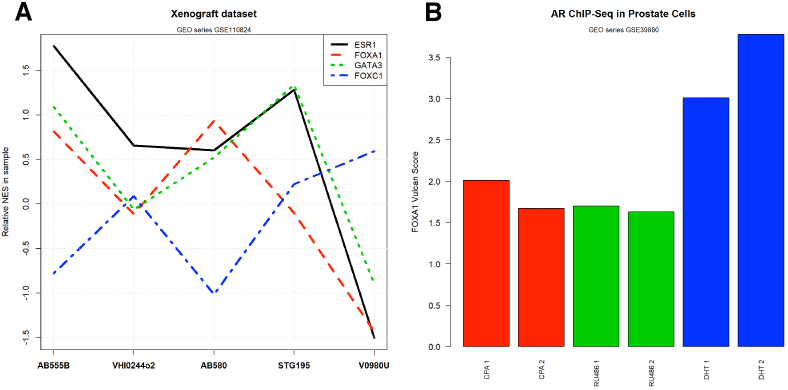
ER occupancy after estradiol treatment in terms of TF network activity. **A**: Global TF network behavior as predicted by VULCAN in our ChIP-seq dataset, highlighting the ESR1 TF at time 0 and 45/90 minutes after estradiol treatment. **B**: Global TF activity after estradiol treatment in MCF7 cells, inferred using the METABRIC network, highlighting TFs significantly upregulated at 45 minutes and 90 minutes. **C**: Global TF activity after estradiol treatment in MCF7 cells, inferred using the METABRIC network, highlighting TFs significantly downregulated at 45 minutes and 90 minutes. **D**: Global TF activity after estradiol treatment in MCF7 cells, inferred using the METABRIC network, highlighting TFs significantly upregulated at 45 minutes but not at 90 minutes. **E**: Global TF activity after estradiol treatment in MCF7 cells, inferred using the METABRIC network, highlighting TFs significantly upregulated at 90 minutes but not at 45 minutes. **F**: Most enriched motif in peaks upregulated at both 45 and 90 minutes after estradiol treatment, as predicted by HOMER.

We performed a Gene Set Enrichment Analysis (GSEA) [16] and an associated Rank Enrichment Analysis (aREA) [15] using the differential binding at gene regulatory regions with time 0 as reference. Individual differential binding signatures for GSEA were calculated using a negative binomial test implemented by DiffBind [18]. The collective contribution of differentially bound sites highlights several ER-related pathways in both the GSEA and aREA analyses [19]–[21] (**Suppl. Fig. S14**). The strongest upregulated GSEA pathway in both time points (**Table S1 and S1**) was derived from RNA-seq in an MCF7 study using estradiol treatment [20], confirming the reproducibility of our dataset.

### VULCAN analysis of ER activation

#### VULCAN identifies coactivators and corepressors of ER

We leveraged the information contained in mutual information networks to establish TF networks enriched in the differential binding patterns induced by estradiol. From our analysis of ER binding, we established four classes of modulation: early coactivators, early corepressors, delayed coactivators and transient coactivators (Fig. 3).

Using VULCAN, we defined TF network activity of occupied regulatory regions (Fig. 3A) according to the binding of ER within their promoter and enhancer regions (limited to 10kb upstream of the Transcription Starting Site to ensure gene specificity). We define early coactivators as those TFs whose network is upregulated at both 45 and 90 minutes (Fig. 3B); these genes include AR, SP1 and CITED1. TFs with opposite behavior (namely TFs whose negative/repressed targets in the ARACNe model are occupied by ER), or “early corepressors”, include GLI4, MYCN and GRHL2 (Fig. 3C). Some TFs appear to have their targets transiently bound at 45 minutes, but then unoccupied at 90 minutes and therefore we dubbed them “transient coactivators’ (Fig. 3D). We further defined TFs active at 90 minutes but not at 45 minutes as “delayed coactivators”, noting these cofactors could be the transient if the response is not completed by 90 minutes. While this category exists, and notably contains both ESR1 and the known ESR1 interactor GATA3, it is just below the significance threshold at 45 minutes (Fig. 3E).

We repeated our TF network activity analysis of ER activation (Fig. 3A-E) on an independent dataset from TCGA and found similar results to those established from the METABRIC-derived network (**Suppl. Fig. S14 to S19**).

To ensure the robustness of the results, we performed a joint analysis of data obtained from both networks. At 45 (Fig. 4A) and 90 minutes (Fig. 4B) we identified candidates, specifically the ESR1, GATA3 and RARA networks, which were consistently and robustly activated by ER in both time points. The joint analysis also identified candidate corepressors, including HSF1 and GRHL2.

**Figure 4:**
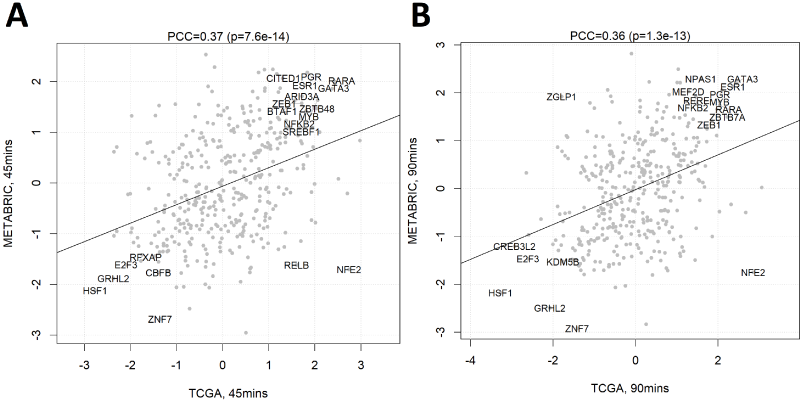
Global TF activity after estradiol treatment using different network models. XY Scatter showing the TF activity as calculated by VULCAN for our differential ChIP-seq analysis of ER binding at 45 minutes (A) and at 90 minutes (B) after stimulation with 100 nM E2. Comparison of results calculated using the METABRIC (y-axis) and TCGA (x-axis) networks shows consistent results know ER interactors including PGR, RARA, GATA3 and GRHL2. GRHL2 activity is notably enriched against. The regulon of ER is also consistently enriched in both networks. Pearson’s Correlation Coefficient (PCC) shown along with significance.

#### VULCAN results are specific to the tissue used for network modeling

Regulatory networks can be tissue-specific due to a variety of biological reasons, such as chromatin status, cofactor availability and lineage-dependent transcriptional rewiring[15]. We tested whether our VULCAN results can be affected by the choice of the ARACNe-inferred regulatory network. In order to do so, we required a gene expression dataset large enough for robust Mutual Information inference (>100 samples), based on the same library preparation and sequencing protocols as the breast cancer TCGA dataset used in our analysis (to remove the possibility of technical differences), but ultimately derived from a tissue as distant as possible from breast cancer (BRCA) on which network models on this study are derived. For this purpose, we computed ARACNe regulatory models on the TCGA dataset for acute myeloid leukemia (AML), a liquid tumor histologically very different from BRCA. This AML-derived network shows globally weaker VULCAN enrichment scores than the BRCA-derived network and a weak positive correlation with the results obtained through breast cancer regulatory models (**Suppl. Fig. S20**). The positive correlation suggests that regulatory networks inferred in breast cancer are tissue specific and can only in part be recapitulated by a leukemia-inferred network.

#### VULCAN is able to predict protein-protein interactions in both patient-derived xenografts (PDX) and prostate cancer

To demonstrate the general applicability of VULCAN, we applied the algorithm to a Breast Cancer Patient-derived xenograft dataset (Gene Expression Omnibus series GSE110824) [22], [23], which showed the expected enrichment of the ESR1, FOXA1 and GATA3 regulons (**Suppl. Fig. S21 and** Fig. 5A) predicting the colocalization of the respective proteins on the chromatin. To further test the generality of VULCAN, we applied the method to another cancer-associated transcription factor type. More specifically, we evaluated an androgen receptor ChIP-seq dataset derived from prostate cancer cell line model LNCaP-1F5 and VCaP (Gene Expression Omnibus Series GSE39880, AR + DHT, RU486 or CPA) [24]. By applying a context specific network built from the TCGA prostate cancer dataset, we could predict functional colocalization of FOXA1 and AR in target genes’ promoters after dihydrotestosterone (DHT) treatment in prostate cell lines (**Suppl. Fig. S22 and** Fig. 5B), validating the known role of FOXA1 in AR-regulated gene transcription in prostate cancer [25][26][27].

**Figure 5:**
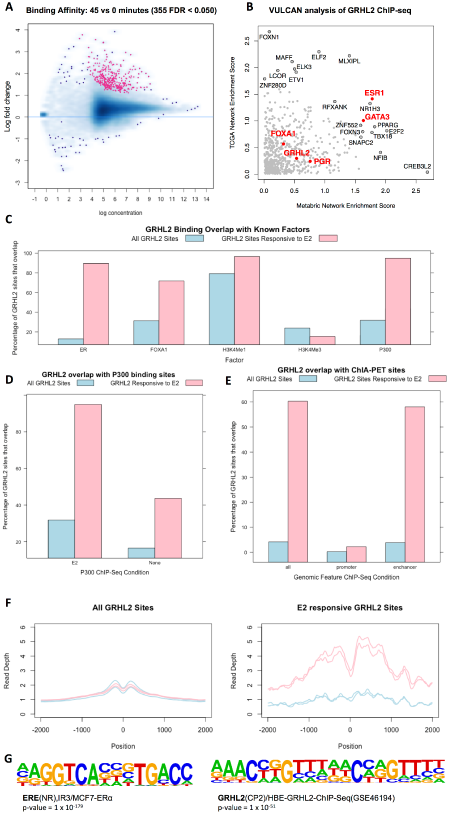
Inferring TF co-occupancy in public datasets with VULCAN. **A.** VULCAN Activity scores for a few TFs derived from the ER-targeted ChIP-seq Breast Cancer Patient-Derived Xenograft (PDX) dataset GSE110824. The behavior of ESR1, FOXA1 and GATA3 is correlated, while FOXC1 shows an inversely correlated pattern (blue line). Interestingly, the sample with the lowest Allred score (V0980U) has the lowest activity and the other luminal markers. **B.** VULCAN Activity scores for FOXA1 in ChIP-seq experiments targeting the Androgen Receptor (AR) in LNCaP-1F5 prostate-derived cells (dataset GSE39880). The bar plots show the relative VULCAN Normalized Enrichment Score calculated on absolute peak intensities after treating cells with Dihydrotestosterone (DHT) and partial AR modulators cyproterone acetate (CPA) and Mifepristone (RU486). FOXA1 network binding is higher in the presence of the strong AR recruiter DHT. This shows an increased FOXA1/AR promoter co-occupancy in DHT-treated cells, in agreement with the conclusions of the study that originated the dataset. Two replicates for each treatment were produced and are reported in matching colors.

#### VULCAN outperforms classical motif analysis

Finally, we compared VULCAN to a classical motif analysis by exploiting the MsigDB C3 collection v6.1 [28] of gene sets, which contain canonical TF-specific binding motifs in their promoters. Our analysis shows the correlation of VULCAN results for two transcription factors (e.g. between GATA3 and ESR1, **Suppl. Fig. S23**) can be relatively high but not significantly overlapping in terms of target genes containing the same canonical motifs (**Suppl. Fig. S24**). We could prove that this non-relationship is general as it extends to the majority of TF-TF pairs that were present in the MsigDB database (**Suppl. Fig. S25**).

#### Optimisation of VULCAN parameters

The network generation algorithm uses established methods to optimise parameters for the RNA-seq input (e.g. ARACNE-AP calculates edge significance based on data-specific permutation test). By default, VULCAN can calculate key settings from the provided ChIP-seq data (e.g. DNA fragment length). Additionally, parameter choice is tunable at the wish of the user.

The distance from promoter TSS can be tuned to the specific organism investigated by the ChIP-seq experiment. In this manuscript, we used 1000nt for Homo sapiens, but it can be lowered to 100nt for bacterial chromosomes, to assign peaks that act as representative for the gene..

### GRHL2 is a novel ER cofactor

In our analysis of ER dynamics, the GRHL2 transcription factor was consistently identified as a key player, using both the METABRIC and TGCA networks. GRHL2 is a transcription factor that is important for maintaining epithelial lineage specificity in multiple tissues [29], [30]. It has previously been predicted to exist in ER-associated enhancer protein complexes [31], but its function in the ER signaling axis is unknown. Therefore, we set out to experimentally validate GRHL2 as an ER cofactor.

There is only a weak, positive correlation between ESR1 and GRHL2 expression in the TCGA and METABRIC breast cancer datasets (**Suppl. Fig. S26**). Furthermore, GRHL2 does not change significantly in different PAM50 subtypes, although it is overexpressed in malignant tissue. The low correlation between GRHL2 expression and subtype implies that the protein is controlled by mechanisms such as phosphorylation [32], subcellular localization or on-chromatin protein-protein interactions.

#### qPLEX-RIME detects a significant increase in the ER-GRHL2 interaction on activation

We undertook a complementary, unbiased, experimental approach combining RIME [9] with TMT [33], called qPLEX-RIME [34], to identify interactors of ER within the ER-chromatin complex. We generated ER qPLEX-RIME data from MCF7 cells treated with estradiol at both 45 and 90 minutes and compared this to the VULCAN dataset (**Suppl. Fig. S27**). We found known ESR1 interactors with both methods, namely HDAC1, NCOA3, GATA3 and RARA. These interactors have positive enrichment according to VULCAN [15], implying the TF’s regulon is over-represented within the differentially bound genes. Importantly, qPLEX-RIME identified a significant increase in the protein-protein interaction between ER and GRHL2 in estrogenic conditions. As GRHL2 has a negative enrichment score in VULCAN, this implies either the protein is recruited by ER to sites that are significantly depleted for GRHL2’s regulon or that GRHL2 is established as having a negative correlation to the genes regulated at these sites, i.e. the protein is a corepressor of the ER.

To assess the chromatin-association of ER and GRHL2, we undertook GRHL2 ChIP-seq in the absence (0 min) or presence (45 min) of E2 (Fig. 6A). VULCAN analysis of the GRHL2 differential binding showed that ER was the key interacting transcription factor, using both the TCGA- and METABRIC-derived networks (Fig. 6B).

**Figure 6:**
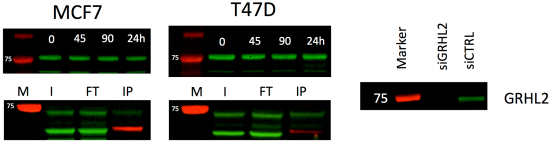
GRHL2 Differential ChIP-seq between 0 and 45 minutes. **A.** Activation of the ER with estro-2-diol results in a genome-wide increase in GRHL2 binding. **B.** VULCAN Analysis of the same data show a significant enrichment for ESR1 sites in both the context of the METABRIC and TGCA networks. The regulon for FOXA1 is also not enriched. Inspection of known FOXA1/GRHL2 sites (e.g. RARa promoter) shows GRHL2 already bound. **C.** Overlap of GRHL2 binding with public datasets shows that E2 responsive GRHL2 sites show considerable overlap with ER, FOXA1 and P300 sites, H3K4Me1 and H3K4Me3 show little enrichment. **D.** Analysis of P300 binding showed a greater overlap of GRHL2 ER responsive sites in the presence of E2 than in control conditions. **E.** Overlap with ER ChIA-PET sites showed enrichment for GRHL2 sites at ER enhancers. **F.** Analysis of Gro-SEQ data (GSE43836) at GRHL2 sites. Blue lines are control samples, pink are samples after stimulation with E2. In general, GRHL2 sites (left) show no change in the levels of transcription on addition of E2; however, E2 responsive GRHL2 sites (right) show a robust increase in transcription on the activation of the ER. **G.** Motif analysis of differentially bound sites gave the top two results as GRHL2 and ER.

We undertook a comparison of GRHL2 binding with public datasets (Fig. 6C). Our analysis showed that GRHL2 sites that responded to estradiol were enriched for ER binding sites (in agreement with our qPLEX-RIME data and VULCAN results) and FOXA1 (compatible with either an ER interaction or the previously reported interaction with MLL3 [31]). Importantly, the changes in GRHL2 binding profiles after E2 treatment were not a result of altered GRHL2 protein levels (Fig. 7). Individual analysis of peaks show that classical ER promoter binding sites, e.g. RARa, were not the target of this redistribution of GRHL2, as these sites were occupied by GRHL2 before E2 stimulation. Motif analysis of the sites within increased GRHL2 occupancy showed enrichment for the full ERE (p-value = 1 × 10^−179^) and the GRHL2 binding motif (p-value = 1 × 10^−51^) (Fig. 6G).

**Figure 7:**
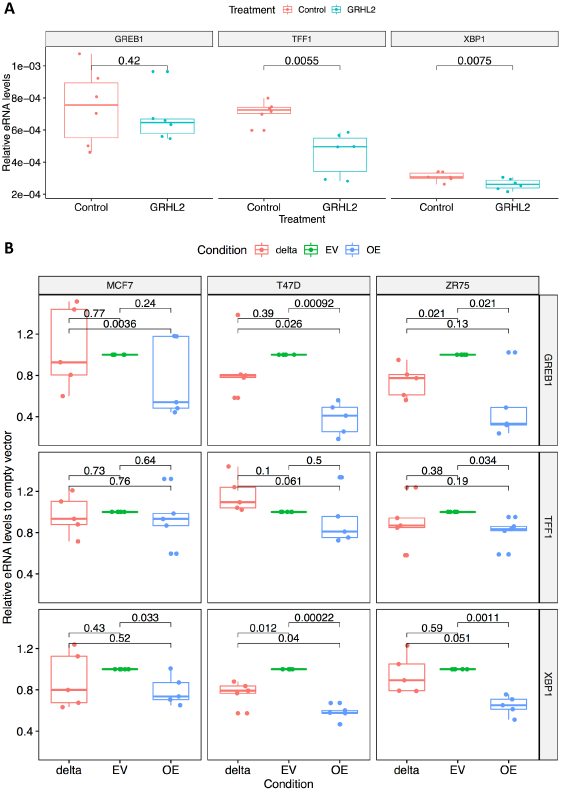
Estrogen Time Course and Co-IP of GRHL2. Analysis by western blot of the GRHL2 showed no changes in the levels of GRHL2 at 45 minutes, 90 minutes or 24 hours after stimulation with estradiol in either MCF7 or T47D. Co-IP of ER (bait, red, Santa Cruz:sc-8002) identified GRHL2 (green, Atlas: HPA004820) as an interactor in estrogenic conditions (M=Marker, I=Input, FT=Flow Through, IP=immunoprecipitation). siRNA knockdown of GRHL2 in MCF7 (right) resulted in loss of the ∼75 kDa band.

To establish if the recruitment of GRHL2 was primarily related to a transcriptional function or the previously described interaction with MLL3, we overlapped our GRHL2 data with that of published H3K4me1/3 [31] and P300 [35] cistromes. While H3K4me occupancy was consistent between conditions, we found P300 binding to be enriched at the E2-responsive GRHL2 sites.

A more detailed analysis of the GRHL2 overlap with P300 sites showed the greatest co-occupancy of GRHL2/P300 sites was when both TFs were stimulated by E2 (Fig. 6D). Moreover, overlap of GRHL2 peaks with ER ChIA-PET data [ENCSR000BZZ] showed that the GRHL2 responsive sites were enriched at enhancers over promoters (Fig. 6E). These findings suggested that the GRHL2-ER complex is involved in transcription at ER enhancer sites.

#### Validation of the ER-GRHL2 interaction by qPLEX-RIME and co-IP

qPLEX-RIME [34] analysis of GRHL2 in both the estrogen-free and estrogenic conditions (Additional File 4) showed high levels of transcription-related protein interactors including HDAC1 (p-value = 6.4 × 10^−9^), TIF1A (p-value = 6.4 × 10^−9^), PRMT (p-value = 6.4 × 10^−9^) and GTF3C2 (p-value = 4.6 × 10^−9^). P-values given for estrogen-free and estrogenic conditions conditions were comparable. The only protein differentially bound to GRHL2 in estrogen-free versus estrogenic conditions was the ER.

We further validated this interaction by co-IP. Our analysis robustly found that GRHL2 and ER interact in both MCF7 and T47D cells (Fig. 7). We further validated the antibody by siRNA knockdown and saw the disappearance of the GRHL2 band at 24 hours.

#### GRHL2 constrains ER binding and activity

We investigated the transcription of enhancer RNAs at these sites using publicly available GRO-seq data [36] [GSE43836] (Fig. 6F). At E2 responsive sites, eRNA transcription was strongly increased by E2 stimulation; by contrast, eRNA transcription was largely independent of E2 stimulation when the entire GRHL2 cistrome was considered. Analysis of a second GRO-seq dataset, GSE45822, corroborated these results **(Suppl. Fig. S28**).

To further explore how GRHL2 regulates ER enhancers, we measured eRNA expression at the GREB1 [37], [38], TFF1 [39]–[41] and XBP1 [42], [43] enhancers after over-expression of GRHL2. At GREB1 and XBP1, increased GRHL2 resulted in reduced eRNA transcription (Fig. 8) (p < 0.05, paired-sample, t-test). Conversely, eRNA production at the TFF1, XBP1 and GREB1 enhancers was moderately increased 24h after GRHL2 knockdown (**Suppl. Fig. S29**). Combining data from all three sites established the effect as significant by paired-sample rank test (p = 0.04, one-tailed paired-sample, wilcoxon test). Collectively, these data demonstrate that GRHL2 constrains specific ER enhancer transcription.

**Figure 8:**
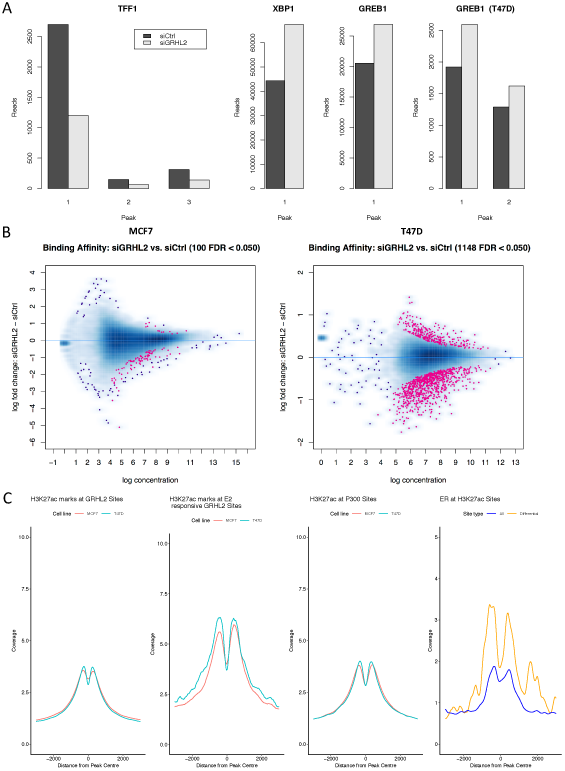
Effect of GRHL2 knockdown after 24 hours on eRNA at E2 responsive binding sites & overexpression of GRHL2 Δ425-437. **A.** Overexpression of GRHL2 in MCF7 resulted in a reduction of eRNA transcribed from the GREB1, TFF1 and XBP1 enhancers. The effect was significant at TFF1 and XBP1 enhancers (p< 0.05, paired t-test). **B.** Overexpression of GRHL2 Δ425-437 (delta) compared to empty vector (EV) and GRHL2 wild-type (OE) at 24 hours. In all 3 cell lines at all 3 loci, overexpression of the wild type (WT) led to a reduction in the mean eRNA production at GREB1, TFF1 and XPB1. This effect was significant in 6 out of 9 experiments (P < 0.05, t-test, one-tailed, paired). Overexpression of GRHL2 Δ425-437 had a reduced effect, that led to a significant reduction in only 2 out of 9 experiments (P < 0.05, t-test, one tailed, paired). Importantly, in 4 out of 9 experiments, WT overexpression had significantly less eRNA production than GRHL2 Δ425-437, suggesting the P300 inhibition domain plays a role in the regulation of eRNA production.

A conserved alpha-helix between residues 425-437 of GRHL2 has previously been shown to inhibit P300 [44]. We therefore overexpressed GRHL2 Δ425-437, a previously demonstrated non-p300-inhibitory mutant [44], [45], in three ER positive breast cancer cell lines (MCF7, T47D and ZR75) and compared levels of eRNA to those recorded for both an empty vector control and for the over-expression of the wild type protein (Fig. 8B). The results of the wild-type study were concordant to those of our previous analysis (Fig. 8A), suggesting in general that overexpression GRHL2 leads to the inhibition of eRNA production at certain ER sites. Importantly, in all cases, the removal of aa 425-437 from GRHL2 led to a reduction in the inhibitory effect than caused by over-expression of the wild-type protein and was found as significant in 5 out of 9 cases test (P < 0.05, t-test, single-tail, paired).

We undertook H3K27ac ChIP-seq after knockdown of GRHL2 by siRNA for 48 hours. In both MCF7 and T47D cells, we saw a significant change in the acetylation marks surrounding GREB1, and in MCF7 we saw an increase at both XBP1 and GREB1 promoters and a decrease at TFF1 (Fig. 9A). Genome-wide, we saw a redistribution of H3K27 acetylation in both cell lines (Fig. 9B). Comparison of the sites altered by GRHL2 knockdown showed a stronger signal for ER binding (Fig. 9C **Right**).

**Figure 9:**
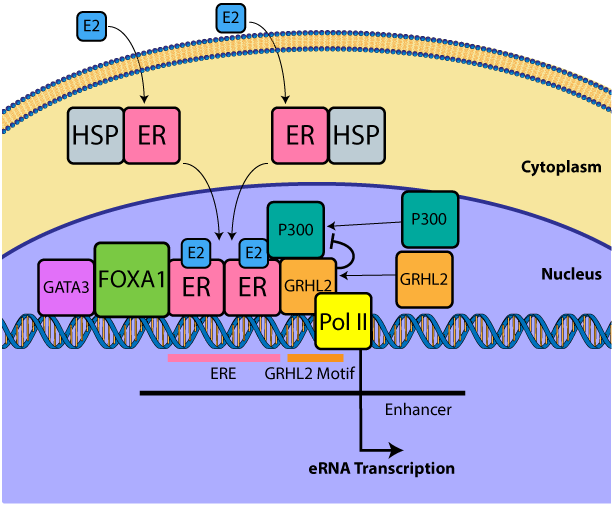
Changes in H3K27ac on knockdown of GRHL2. **A.** The effect of silencing GRHL2 on H3K27ac at 48 hours in MCF7 and T47D cell lines was monitored by ChIP-seq. Analysis of sites proximal to TFF1, XBP1 and GREB1 showed significant changes in acetylation at all three sites in MCF7. Significant changes were only found at GREB1 in T47D (top right). While XBP1 and GREB1 show an increase in histone acetylation on silencing GRHL2, TFF1 showed the reverse effect. **B.** Genomewide, the effects of silencing GRHL2 led to significant redistribution of H3K27ac in both the MCF7 and T47D cell lines, with both showing an increase and decrease in the histone mark dependent on site. **C.** From left to right. Coverage as calculated by Homer. H3K27ac was found at GRHL2 sites in both MCF7 and T47D cells, in particular at the E2 responsive sites. The same mark was also found at P300 sites as expected. Analysis of ER binding at H3K27ac sites showed an enrichment for ER binding at the H3K27ac sites that were most responsive to knockdown of GRHL2 in MCF7 cells.

## Discussion

### VirtUaL ChIP-seq Analysis through Networks – VULCAN

VULCAN is valuable for the discovery of transcription factors acting as coregulators within chromatin-bound complexes that would otherwise remain hidden. The challenge of highlighting cofactors from a ChIP-seq experiment lays in the infeasibility of reliable proteomic characterization of DNA-bound complexes at specific regions. On the other hand, while RNA-seq is arguably the most efficient technique to obtain genome-wide quantitative measurements, any transcriptomic approach cannot provide a full picture of cellular responses for stimuli that are provided on a shorter timescale than mRNA synthesis speed, such as the estradiol administration described in our study. VULCAN, by combining RNA-seq derived networks and ChIP-seq cistrome data, aims at overcoming limitations of both. Most notably, our method can work in scenarios where candidate cofactors do not have a well-characterized binding site or do not even bind DNA directly.

Through comparative analysis we have robustly shown that VULCAN is able to outperform other readily available methods for the prediction of on-chromatin interactions of transcription factors. VULCAN achieves this through the integration of ChIP-seq and tumour transcriptional data. The inherent limitation of our method therefore is that tumour transcriptional data must be available in sufficient quantity to build the underlying network for the analysis, whereas tools based on predefined networks have no such limitation. In the majority of cases this is not a challenge as projects like the TCGA provide transcriptome-wide data for a range of cancers. It is therefore only in the cases of rarer disease types (such as Neuroendocrine Tumor) and orphan tissues that this limitation will be problematic as these are poorly represented in public data. Even so in these cases where specific networks cannot be generated, pan-tissue regulatory networks are currently being developed to overcome this limitation and these could be adapted for VULCAN. [46]

By developing VULCAN, we have been able to rediscover known cofactors of the estradiol-responsive ER complex and predict and experimentally validate a novel protein-protein interaction.

### GRHL2-ER Interaction

#### GRHL2 has a key role in regulating EMT

In the 4T1 tumor model, GRHL2 was found to be significantly downregulated in cells that had undergone EMT [29]. The same study showed that knockdown of GRHL2 in MCF10A – an ER-negative cell line – led to loss of epithelial morphology. Overall, this suggested that the GRHL2 transcription factor plays an essential role in maintaining the epithelial phenotype of breast cells. Similar results were observed with the MDA-MB-231 model, where expression of GRHL2 resulted in reversal of EMT [30]. This result has been recapitulated in hepatocytes, where GRHL2 was found to suppress EMT by inhibiting P300 [44]. The ability to suppress EMT has also been noted in prostate cancer, another cancer driven by a steroid hormone receptor (AR), and the genes regulated by GRHL2 are linked to disease progression [47].

#### GRHL2 a novel corepressor of ER eRNA production

These earlier data combined with the link between GRHL2 expression and patient survival indicate a significant role for GRHL2 in the progression of breast cancer. However, its role in the ER signaling axis has, until now, been unknown. Here, we show that GRHL2 performs its activity at a subset of ER enhancers. Overexpression of GRHL2 resulted in a significant decrease in eRNA production at the TFF1 and XBP1 enhancers and, in agreement with previous studies that correlate eRNA transcription with gene expression [48]–[50], we found the measured eRNA decrease was concurrent with a significant downregulation in the expression of the corresponding gene.

These results are consistent with previous findings that GRHL2 inhibits P300 [44] and, while the ER complex results in the activation of eRNA transcription at these sites, that GRHL2 plays a role in fine-tuning or modulating this process.

#### GRHL2 role in the ER signaling axis is independent to its role in tethering MLL3

In breast cancer, GRHL2 has previously been shown to directly interact with FOXA1, which may contribute to tethering of the histone methyltransferase MLL3 and, consequently, epigenetic marks at GRHL2/FOXA1 binding sites [31]. Our analysis, however, showed no particular enrichment for H3K4me1/3 marks at E2 responsive GRHL2 sites compared to other GRHL2 binding sites and our proteomic analysis of interactors showed a strong association with proteins related to transcription. We proposed that these ER responsive sites are related to a role of GRHL2 in a transcriptional process independent of its interaction with MLL3. This was supported by evidence of a significant overlap with binding of the coactivator P300, transcriptional proteins detected by qPLEX-RIME analysis of GR, and a pronounced increase in eRNA transcription at E2 responsive GRHL2 sites.

#### ER is bound more strongly at active enhancers (H3K27ac) that are altered by siGRHL2

Knockdown of GRHL2 led to a genome-wide remodeling of H3K27ac marks, found at active enhancers, confirming a role of GRHL2 in partially regulating these sites. Detailed inspection of the data showed a significant increase of these marks around the XBP1 and GREB1 genes, supporting our hypothesis that GRHL2 has a partial inhibitory role within the ER regulon. The result was further supported by finding enrichment of ER binding events at H3K27ac marks altered by GRHL2 knockdown (Fig. 9C, **Right Panel**). The more complex effect on H3K12ac, when compared to the effects on eRNA production, is likely a result of the diversity of roles that GRHL2 holds within the cell, leading to a host of downstream effects in the regulation of chromatin recruitment of key factors such as MLL3, ER and FOXA1[31]

#### Deletion of the P300 inhibitory α-helix from GRHL2 reduces the protein’s ability to repress the production of eRNA at ER bound enhancer sites

Finally, to clarify if inhibition of P300 was occurring, we generated and overexpressed GRHL2 lacking the inhibitory alpha-helix at between amino acids 425-437. In all cases, GRHL2 Δ425-437 had reduced an inhibitory effect compared to overexpression of the wild-type, confirming that GRHL2 primarily plays a repressive role at these sites.

## Conclusion

VULCAN is built on state-of-the-art network analysis tools previously applied to RNA-seq data. By adapting network-based strategies to ChIP-seq data, we have been able to reveal novel information regarding the regulation of breast cancer in a model system.

We have demonstrated that the VULCAN algorithm can be applied generally to ChIP-seq for the identification of new key regulator interactions. Our method provides a novel approach to investigate chromatin occupancy of cofactors that are too transient or for which no reliable antibody is available for direct ChIP-seq analysis.

Further, because of our use of clinical data, VULCAN results are both more likely to be relevant and are specific to the disease type studied, as demonstrated in the loss of signal when using a control coexpression network generated from an alternative disease type.

VULCAN enabled us to identify the GRHL2-ER interaction and that GRHL2 plays a repressive role. Further analysis showed the process to be independent to the previously reported interaction with FOXA1 and MLL3 [31]. Our conclusion, therefore, is that GRHL2 has a second, previously undescribed role that regulates transcription at specific estrogen responsive enhancers (**Fig. 10**).

**Figure 10:** Overview of the role of GRHL2 in ER activation. On activation of the ER by the ligand E2, the protein is released from a complex containing HSPs and translocates to the nucleus. The holoER dimer forms a core complex at Estrogen Response Elements (ERE) with FOXA1 (pioneer factor) and GATA3. ER further recruits P300 and GRHL2. GRHL2 has an inhibitory effect on P300 (a transcriptional activator interacting with TFIID, TFIIB, and RNAPII), thereby reducing the level of eRNA transcription at enhancer sites. Overexpression of GRHL2 further suppresses transcription, while knockdown of GRHL2 reverses the process.

Given the central role of the ER in breast cancer development and GRHL2’s own ability to regulate EMT, the discovery that ER recruits GRHL2 leading to the altered eRNA production is an important step in enhancing our understanding of breast cancer and tumorigenesis.

## Methods

### VULCAN

An implementation of VULCAN in R is available on Bioconductor.org [https://bioconductor.org/packages/release/bioc/html/vulcan.html] and the scripts to replicate our analysis are available as Rmarkdown files. Unless otherwise specified, all p-values were Bonferroni-corrected.

### Sample preparation

MCF7 cells were obtained from the CRUK Cambridge Institute collection, authenticated by STR genotyping and confirmed free of mycoplasma. All cells were maintained at 37 °C, 5% CO_2_. For each individual ChIP pull-down, we cultured 8 × 10^7^ MCF7 cells (ATCC) across four 15 cm diameter plates in DMEM with 10% FBS, Glutamine and Penicillin/Streptomycin (Glibco). Five days before the experiment, the cells were washed with phosphate buffered saline (PBS) and the media was replaced with clear DMEM supplemented with charcoal treated serum. The media was refreshed every 24 hours, which halted the growth of the cells and ensured that majority ER within the cell was not active. On day 5, the cells were treated with estradiol (100 nM). At the appropriate time point, the cells were washed with ice cold PBS twice and then fixed by incubating with 10mL per plate of 1% formaldehyde in unsupplemented clear media for 10 minutes. The reaction was stopped by the addition of 1.5mL of 2.5 M glycine and the plates were washed twice with ice cold PBS. Each plate was then scraped in 1 mL of PBS with protease inhibitors (PI) into a 1.5 mL microcentrifuge tube. The cells were centrifuged at 8000 rpm for 3 minutes at 4 °C and the supernatant removed. The process was repeated for a second wash in 1 mL PBS+PI and the PBS removed before storing at −80 °C.

### ChIP-seq

Frozen samples were processed using established ChIP protocols [51] to obtain DNA fragments of ∼300 bp in length. The libraries were prepared from the purified DNA using a Thruplex DNA-seq kit (Rubicon Genomics) and sequenced on the Illumina HiSeq Platform. Sequencing data is available from Gene Expression Omnibus, accession GSE109820 & GSE123475.

### Differential binding analysis

Sequencing data was aligned using BWA [52] to the human genome (hg19). Reads from within the DAC Blacklisted Regions was removed before peak calling with MACS 2.1 [53] on default parameters. The aligned reads and associated peak files were then analyzed using DiffBind [18] to identify significant changes in ER binding.

### Gene Set Enrichment Analysis (GSEA)

Gene Set Enrichment Analysis (GSEA) was performed as described by Subramanian et al. [54] using the curated pathway collection (C2) from MSIGDB v 5.0 with 1000 set permutations for each pathway investigated, followed by Benjamini Hochberg P-value correction.

### Motif analysis

Motif analysis of the binding regions was undertaken with Homer v4.4 [55] using default parameters. Motif logo rendering was performed using Weblogo v2.8.2 [56]

### VULCAN analysis

We reconstructed a regulatory gene network using ARACNe-AP as described by Alvarez [57]. RNA-seq breast cancer data was downloaded from TCGA on January 2015 and VST-Normalized as described by Anders and Huber [58]. The ARACNe transcriptional regulation network was imported into R using the VIPER BioConductor package and it was interrogated using the differential binding profiles from our ChIP-seq experiment as signatures, 45 minutes vs control and 90 minutes vs control. The peak-to-promoter assignment was performed using a 10kb window with respect to the transcription starting site (TSS) of every gene on the hg19 human genome. The algorithm msVIPER (multi-sample Virtual Inference of Protein activity by Enriched Regulon analysis) was then applied, leveraging the full set of eight replicates per group, with 1000 signature permutations and default parameters.

## qPLEX-RIME

Samples were prepared as previously described for RIME [9], protocol was modified to include TMT isobaric labels for quantification [34].

## TF Binding Overlap

Publicly available data was downloaded as described in the source publication [3], [31], [35], [36] and overlap was calculated with bedtools (v2.25.0). Presented data was normalized as a percentage of GRHL2 sites.

## eRNA quantification

MCF7 cells were transfected with Smart Pool siRNA (Dharmacon, L-014515-02), siControl, GRHL2 overexpression vector (Origene, RC214498), GRHL2 Δ425-437 (Origene), or empty control vector using Lipofectamine 3000 (Thermo Fisher Scientific) according to the manufacturer’s protocol in 6-well format. Expression was monitored by rtPCR using TaqMan assay with GAPDH as a control transcript. Knockdown efficiency was ∼75% and the GRHL2 overexpression vector led a 730-fold increase in expression over control plasmid. 1 ug of purified RNA was reverse transcribed with Superscript III reverse transcriptase (Thermo Fisher Scientific, 18080085) using random primers (Promega, C1181) according to manufacturer instructions. eRNAs were quantified with qPCR using Power SYBR™ Green PCR Master Mix (Thermo Fisher Scientific, 4367660) and denoted as relative eRNA levels after normalizing with UBC mRNA levels.

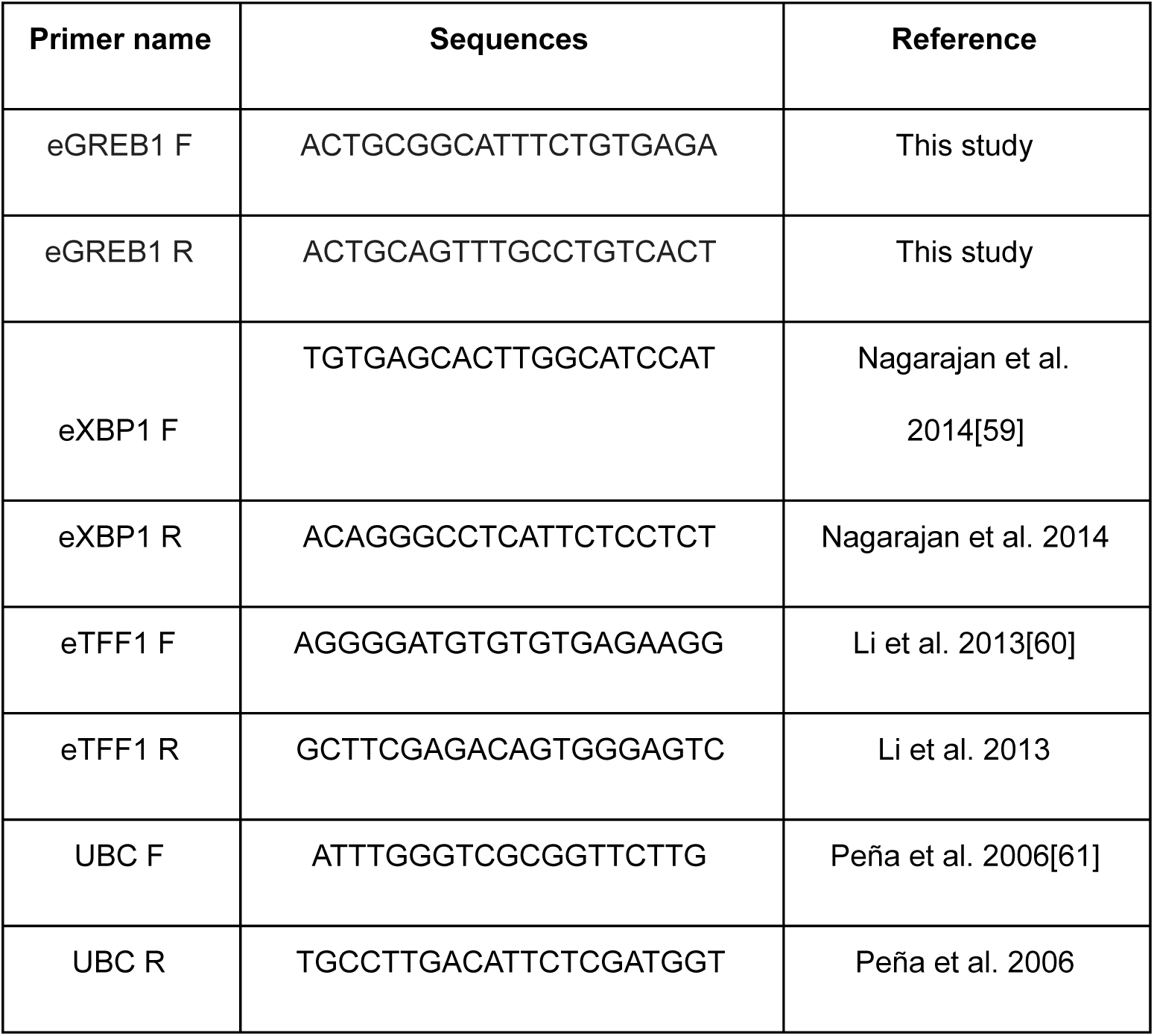

## Co-Immunoprecipitation

ERα (F10) antibody, SantaCruz (sc-8002) was cleaned using Amicon 10K Buffer Exchange Column (EMD, Cat # UFC501096) to remove the Sodium Azide. 2.5 μ g ERα (F10) antibody rotated overnight at 4°C with 100μl Dynabeads Protein A, Invitrogen (10001D).

Nuclear lysate was harvested via cell lysis (20mM Tris-HCl, 20mM NaCl, 0.2mM EDTA) followed by nuclear lysis (20mM Tris-HCl, 20mM NaCl, 0.2mM EDTA 1% Igepal). Nuclear lysate was then incubated overnight at 4°C with the Dynabeads Protein A. Elution via 10 minutes incubation at 70°C with 1X NuPAGE LDS Sample Buffer, Invitrogen (NP0007) and 1X NuPAGE Sample Reducing Agent, Invitrogen (NP0004) and subjected to Western Blotting.

## Knockdown of GRHL2

Knockdown of GRHL2 was undertaken using ON-TARGETplus SMARTpool Human GRHL2, Dharmacon (#L-014515-02-0050) and Lipofectamine® RNAiMAX Reagent Protocol (Thermo) according to the manufacturer’s protocol. Control samples were prepared following the same method using ON-TARGETplus Control pool Non targeting pool, Dharmacon (#D-001810-10-50) in place of siGRHL2.

## Declarations

### Author Contributions

ANH, FMG & LAS conceived and designed the experiments, ANH undertook ER experimentation, ANH & FMG undertook the analysis of ER data, ANH & FMG designed VULCAN, FMG implemented the VULCAN analysis, AEC, ANH & AD undertook GRHL2 experimentation and SN undertook eRNA quantitation. ANH, AEC & LAS undertook analysis of GRHL2 data. ANH, FMG, AEC & LAS wrote the manuscript.

#### Acknowledgements

We would like to acknowledge the support of the University of Cambridge, Cancer Research UK and Hutchison Whampoa Limited.

We are grateful for the support of Eva Papachristou in undertaking the qPLEX-RIME analysis of our samples in advance of the her initial publication describing the method.

We would like to acknowledge the contribution from the CRUK Genomics, Proteomics and Bioinformatics core facilities in supporting this work.

### Ethics Approval and Consent to Participate

Not applicable.

### Availability of Data and Materials

All sequence data are publically available at the GEO, accession numbers GSE109820 [62] and GSE123475 [63]. Data was from previously published datasets were also acquired from GEO with accession numbers GSE110824 [64], GSE39880 [65], GSE45822 [66], GSE43836 [67].

Code for data analysis is provided as a Rmarkdown document and supporting data is available from https://github.com/andrewholding/VULCANSupplementary [68]. For convenience, we have provided VULCAN as a BioConductor package https://bioconductor.org/packages/release/bioc/html/vulcan.html [69] along with a supporting data package https://bioconductor.org/packages/release/data/experiment/html/vulcandata.html [70].

The markdown code [68] created to support this research is distributed under GPLv3. The VULCAN package [69] is distributed under LGPLv3.

## Abbreviations

AR: Androgen Receptor
ARACNe-AP: AccuRate Algorithm for reConstruction of Network through Adaptive Partitioning
CCLE: Cancer Cell Line Encyclopedia
ChIP: Chromatin ImmunoPrecipitation
E2: Estradiol
ER: Estrogen Receptor-Alpha
ERE: Estrogen Response Elements
eRNA: Enhancer RNA
GSEA: Gene Set Enrichment Analysis
GRHL2: Grainyhead Like Transcription Factor 2
METABRIC: MolEcular TAxonomy of BReast cancer International Consortium PBS Phosphate Buffered Saline
PI: Protease Inhibitors
PR: Progesterone Receptor
TCGA: The Cancer Genome Atlas
TF: Transcription Factor
VIPER: Virtual Inference of Protein activity by Enriched Regulon analysis VULCAN VirtUaL ChIP-seq Analysis through Networks

