## Supplementary Figures for "VULCAN integrates ChIP-seq with patient-derived co-expression networks to identify GRHL2 as a key co-regulator of ERa at enhancers in breast cancer"

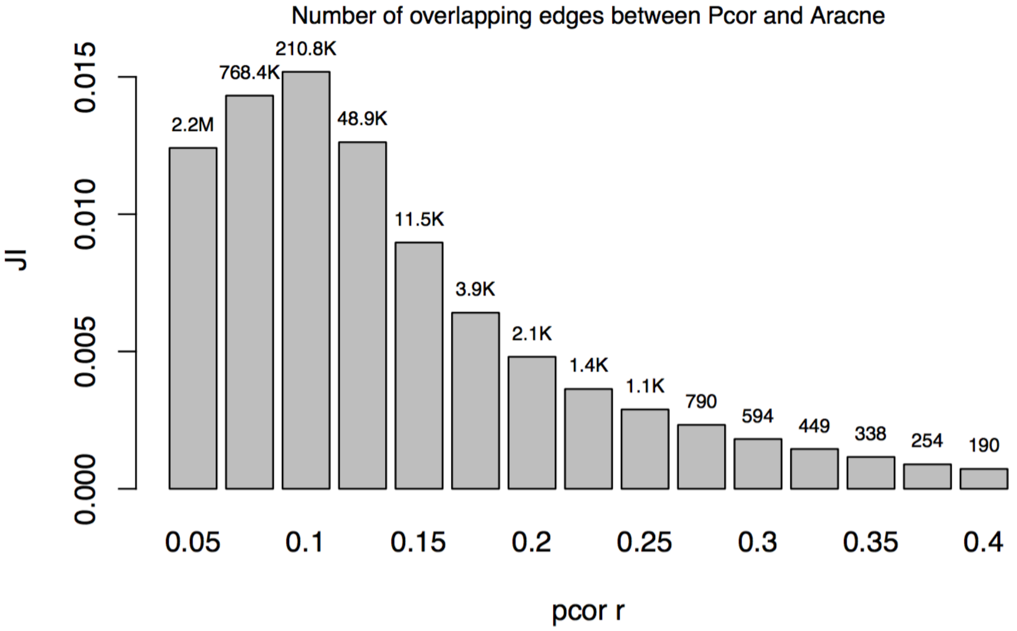


**Fig. S1: Sheer number of overlapping edges at different Pcor thresholds.**

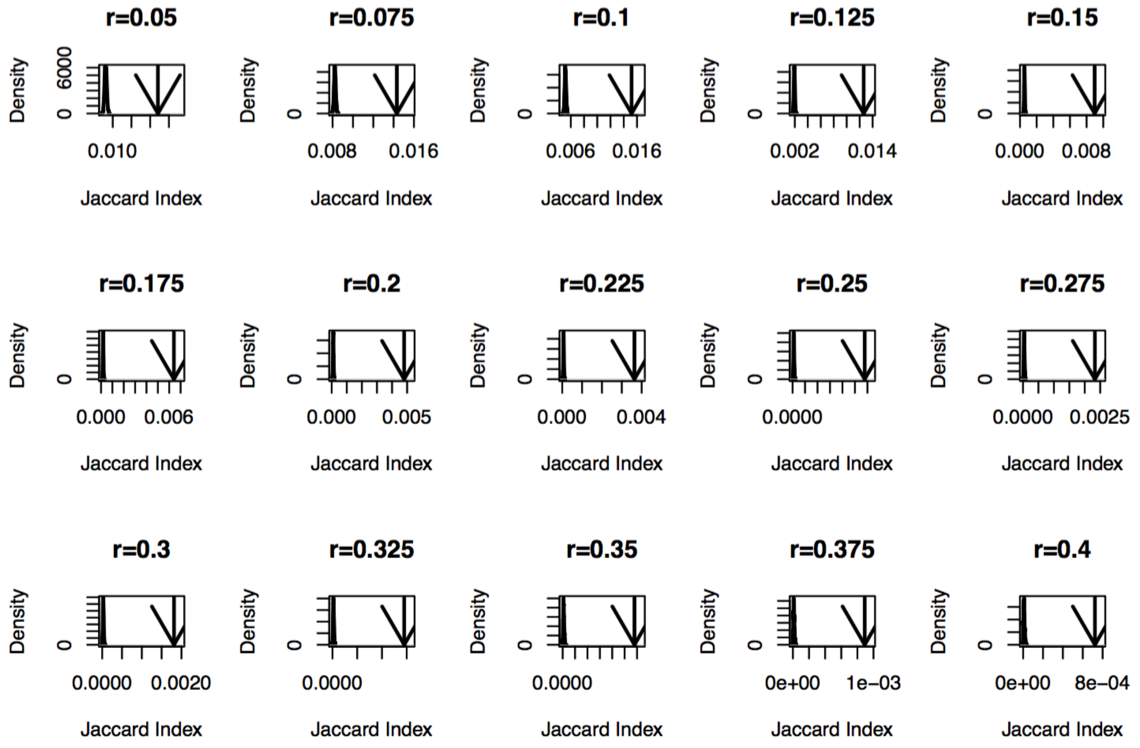


**Fig. S2: Comparing the Jaccard Index between the ARACNe network and Partial Correlation networks (indicated as arrows) and Random networks (indicated as distributions).**

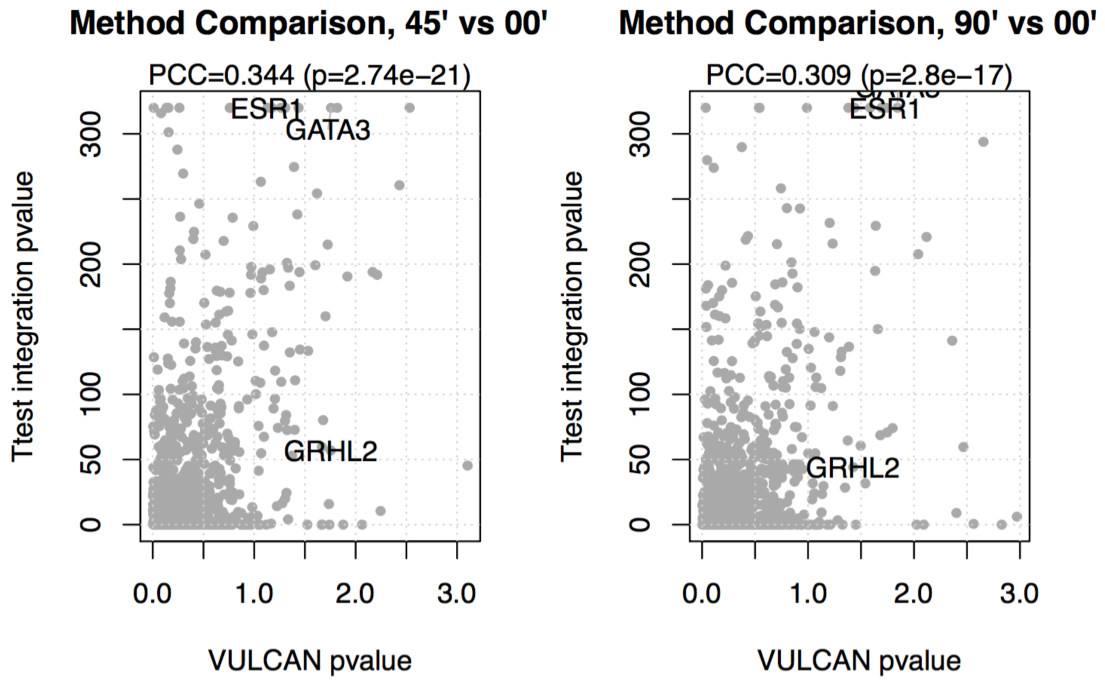


**Fig. S3: Comparison between VULCAN/VIPER and T-test integration.**

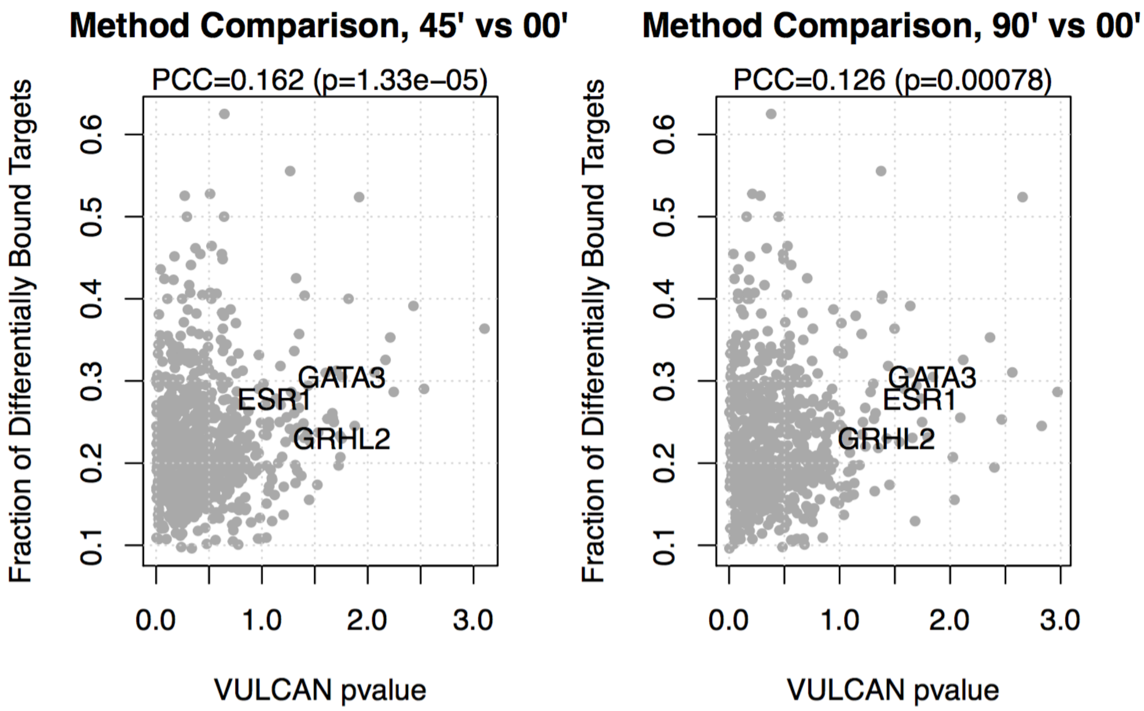


**Fig. S4: Comparison between VULCAN/VIPER and a fraction of targets found method.**

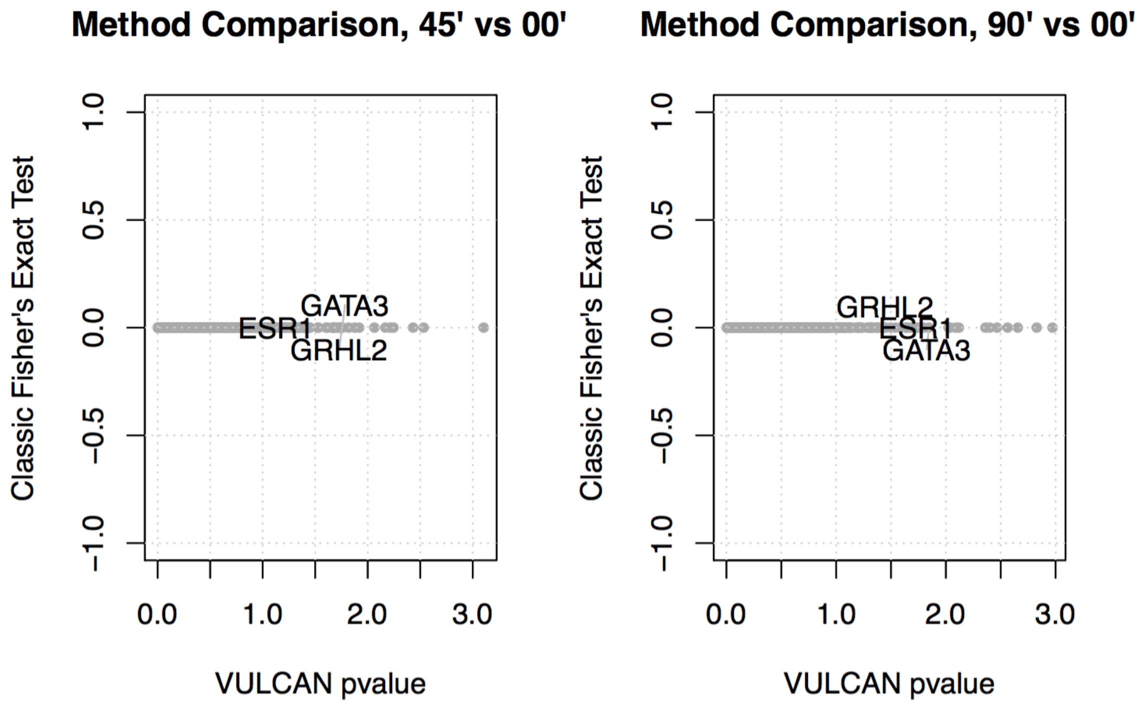


**Fig. S5: Comparison between VULCAN/VIPER and Fisher's Exact Test method.**

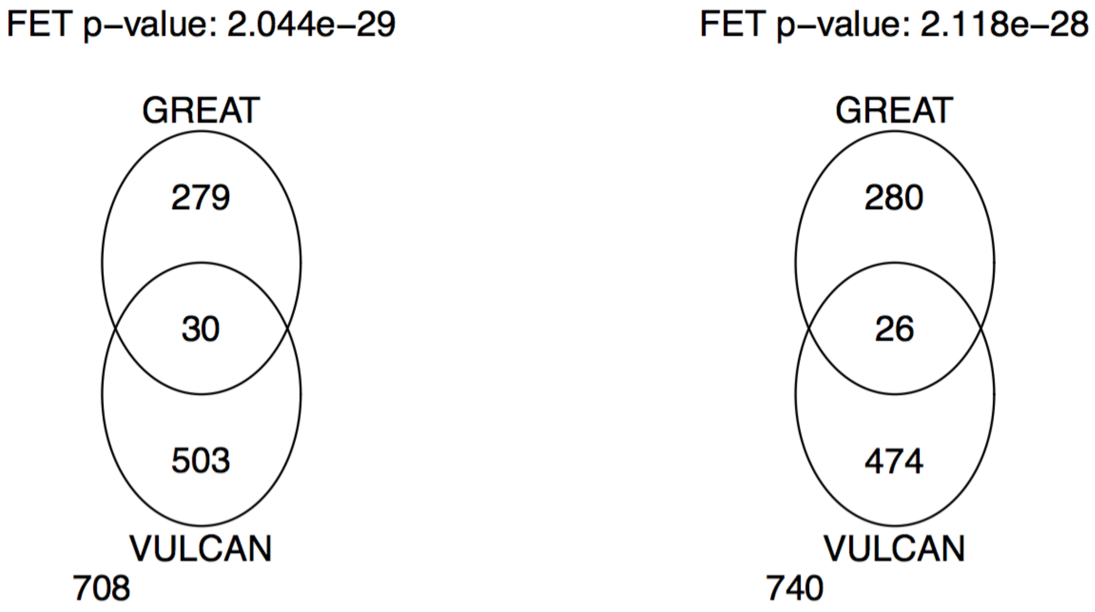


**Fig. S6: Comparison of results from the VULCAN and GREAT methods.**

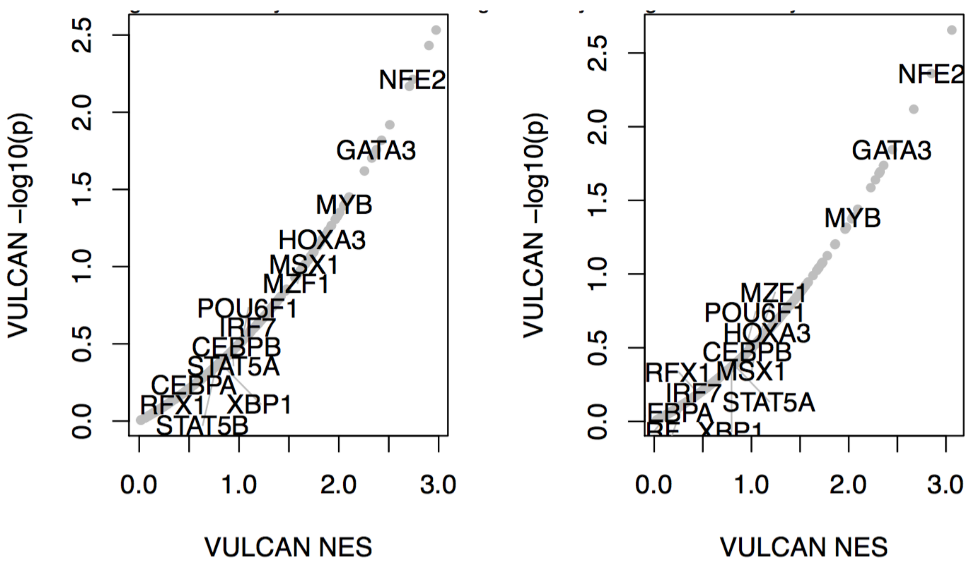


**Fig. S7: Comparison of results from the VULCAN and ChIP-enrich methods.**

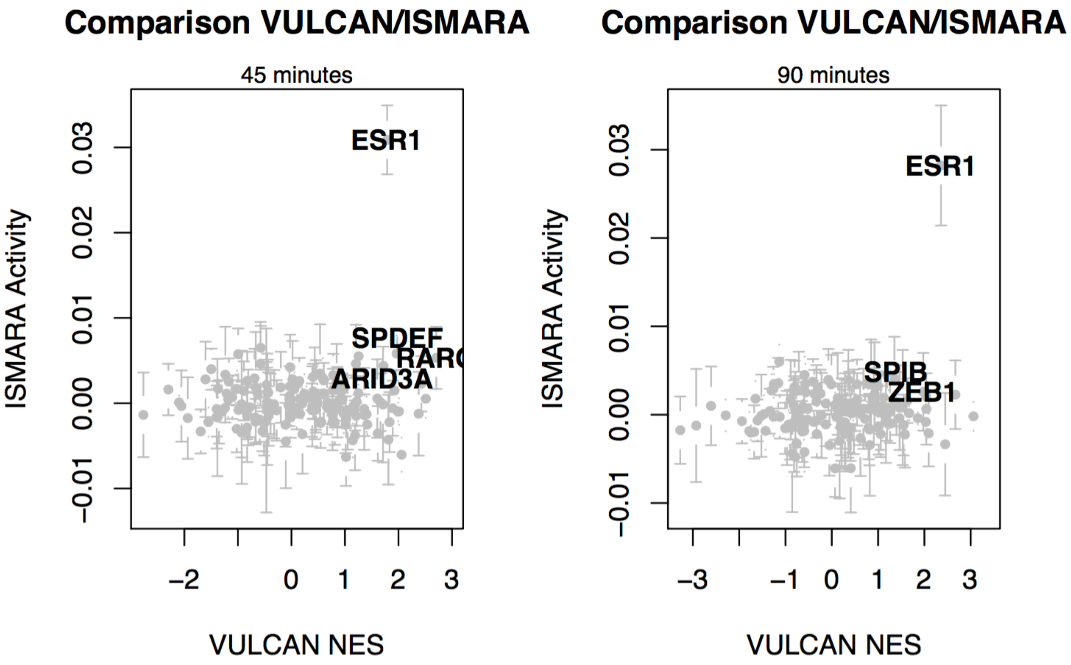


**Fig. S8: Comparison of results from the VULCAN and ISMARA methods.**

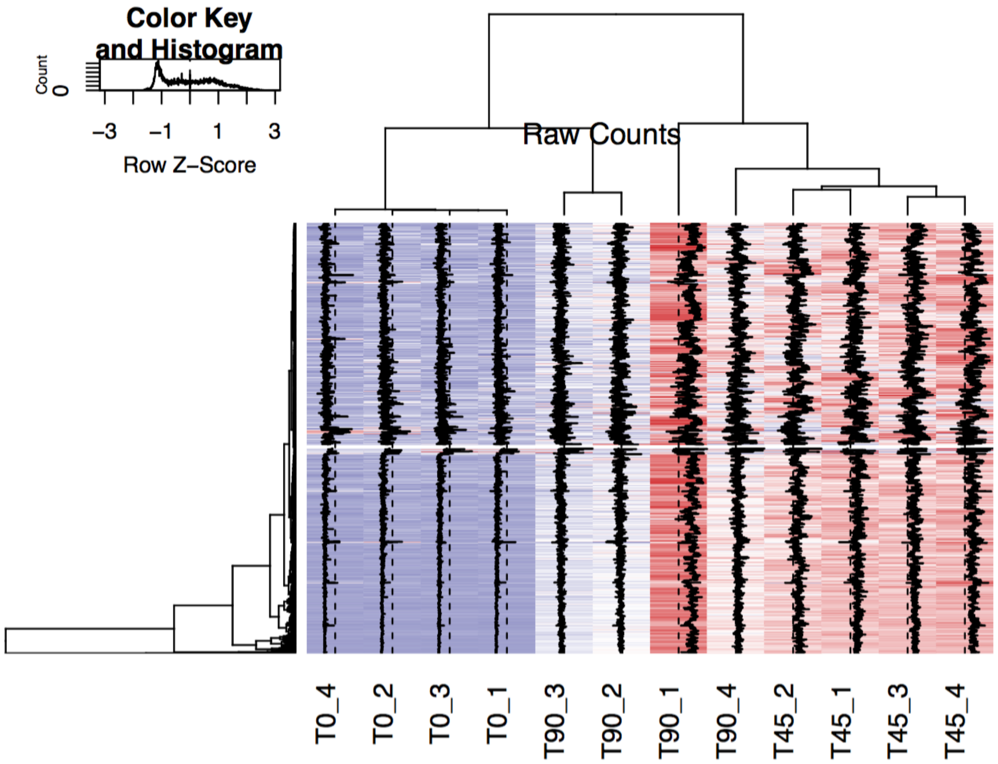


**Fig. S9: Dataset clustering with Raw Counts.**

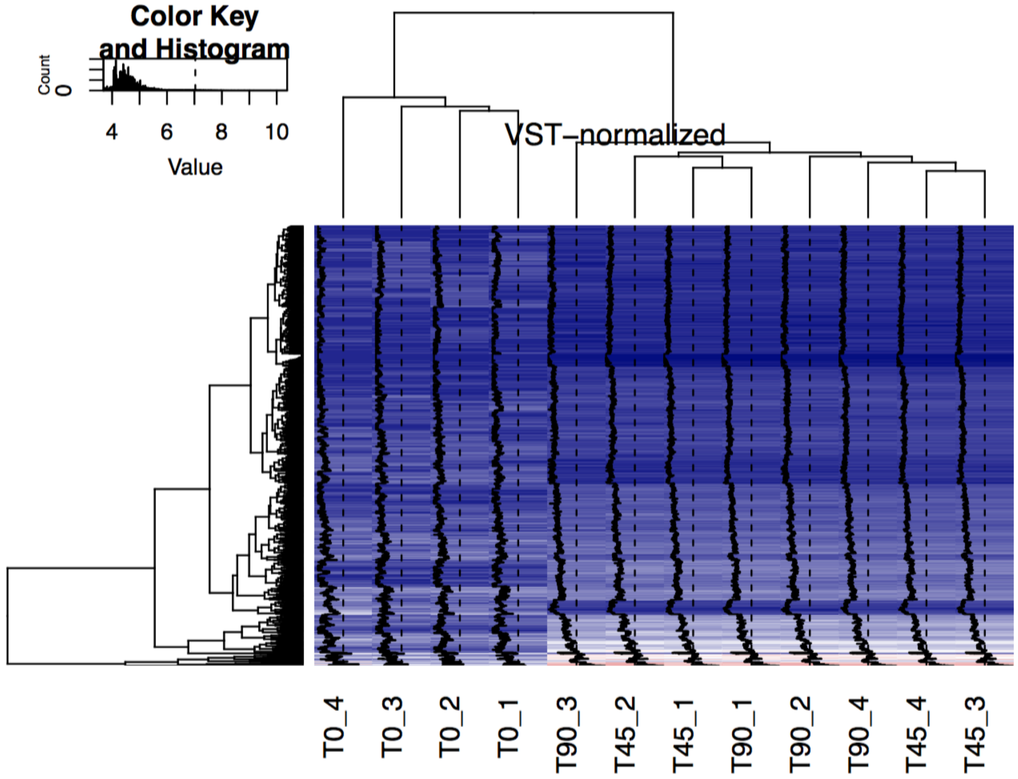


**Fig. S10: Dataset clustering with VST-normalized Counts.**

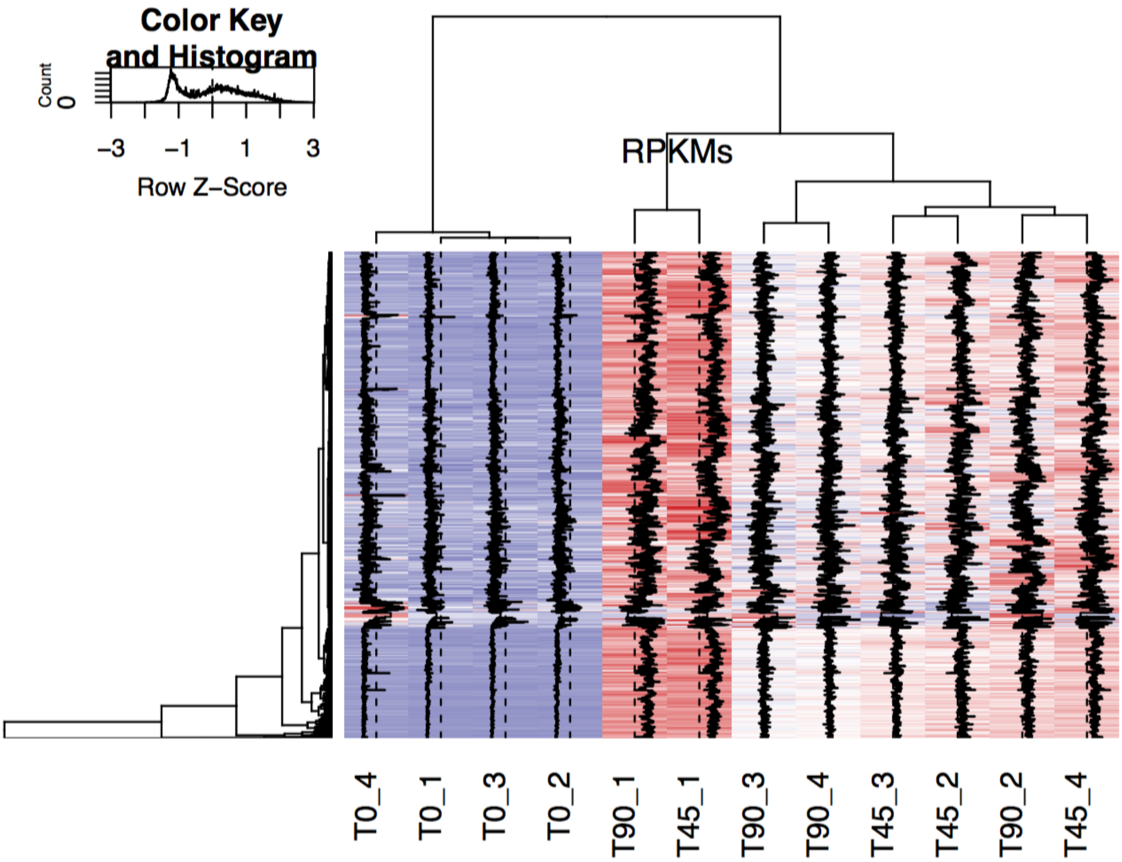


**Fig. S11: Dataset clustering with RPKMs.**

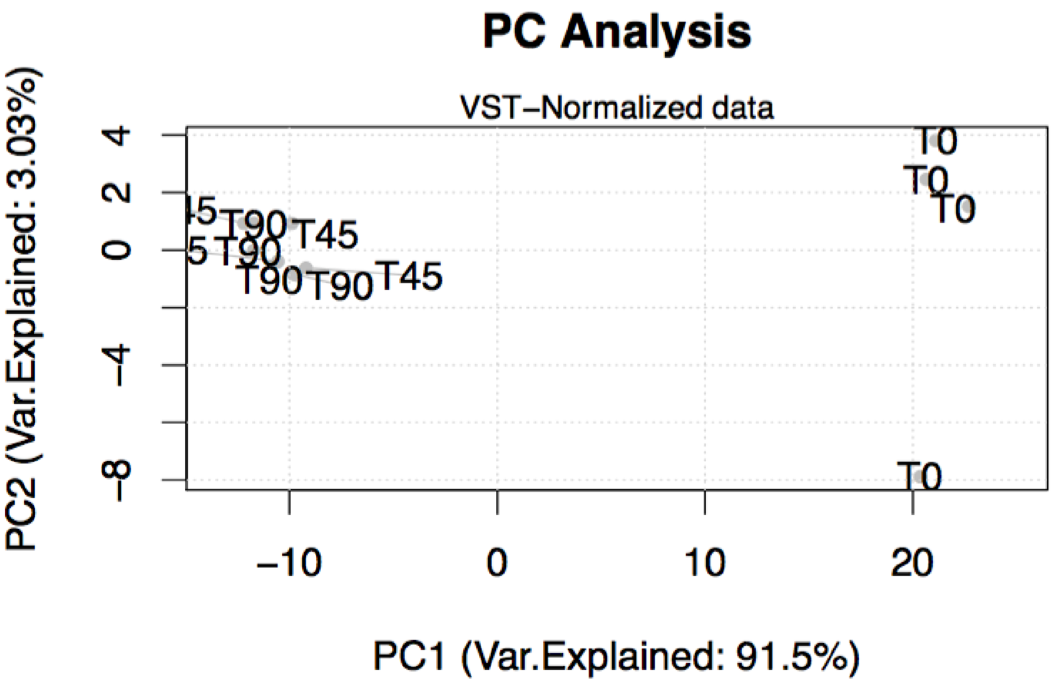


**Fig. S12: Principal Component Analysis of the dataset, highlighting components 1 and 2.**

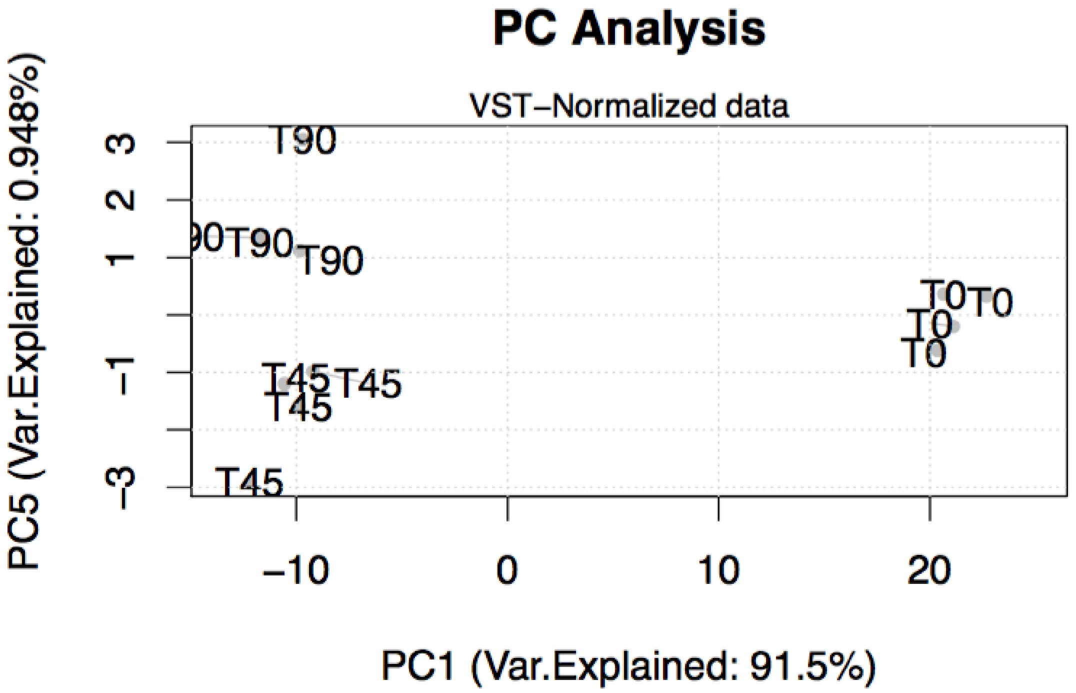


**Fig. S13: Principal Component Analysis of the dataset, highlighting components 1 and 5.**

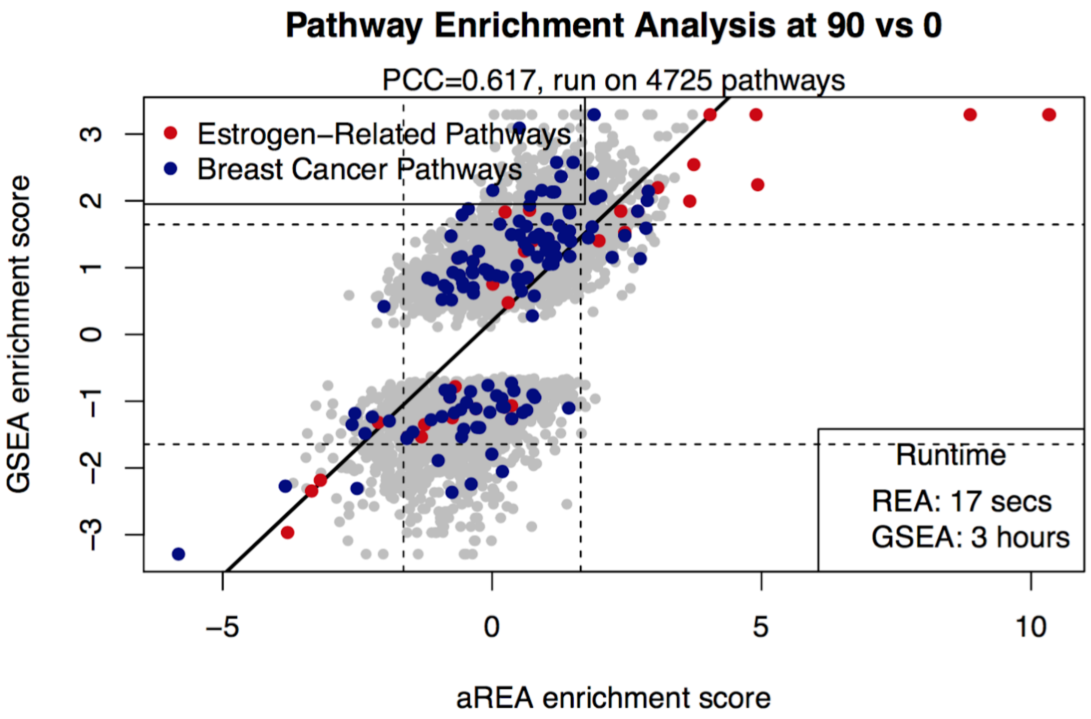


**Fig. S14: Comparison of GSEA and aREA on a differential binding signature.**

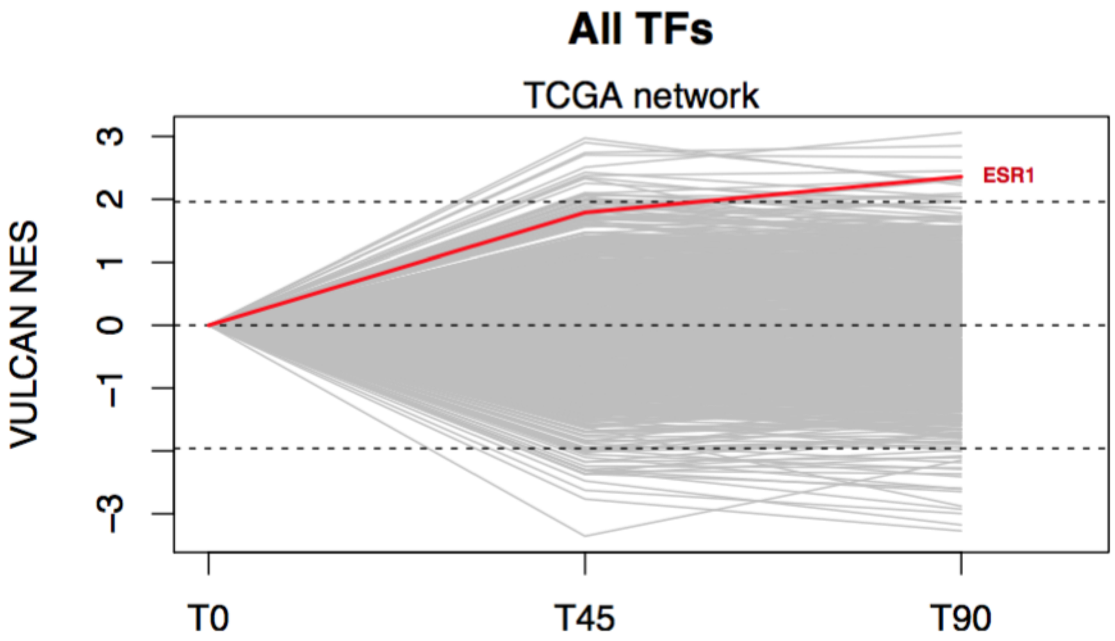


**Fig. S15: Global TF activity after Estradiol Treatment in MCF7 cells, inferred using the TCGA network, highlighting the ESR1 TF as an example.**

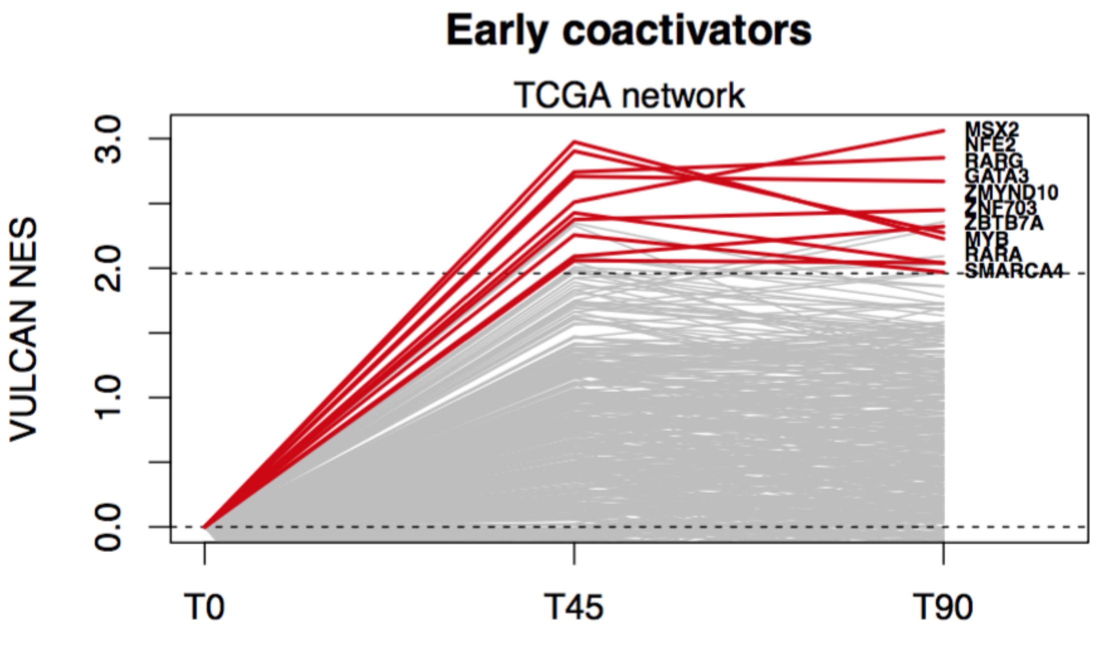


**Fig. S16: Global TF activity after Estradiol Treatment in MCF7 cells, inferred using the TCGA network, highlighting TFs significantly upregulated at 45 minutes and 90 minutes.**

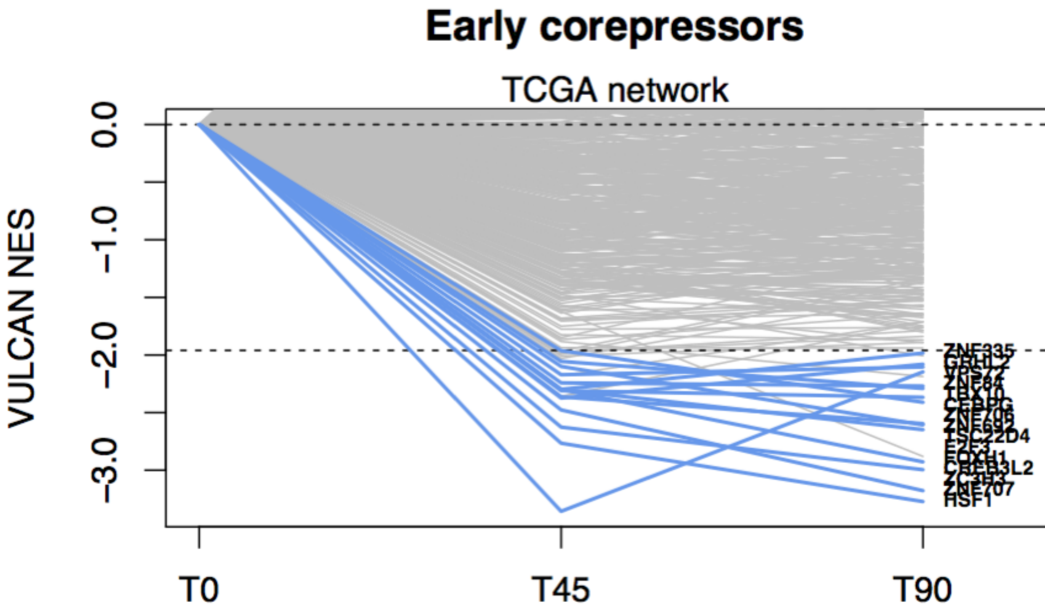


**Fig. S17: Global TF activity after Estradiol Treatment in MCF7 cells, inferred using the TCGA network, highlighting TFs significantly downregulated at 45 minutes and 90 minutes.**

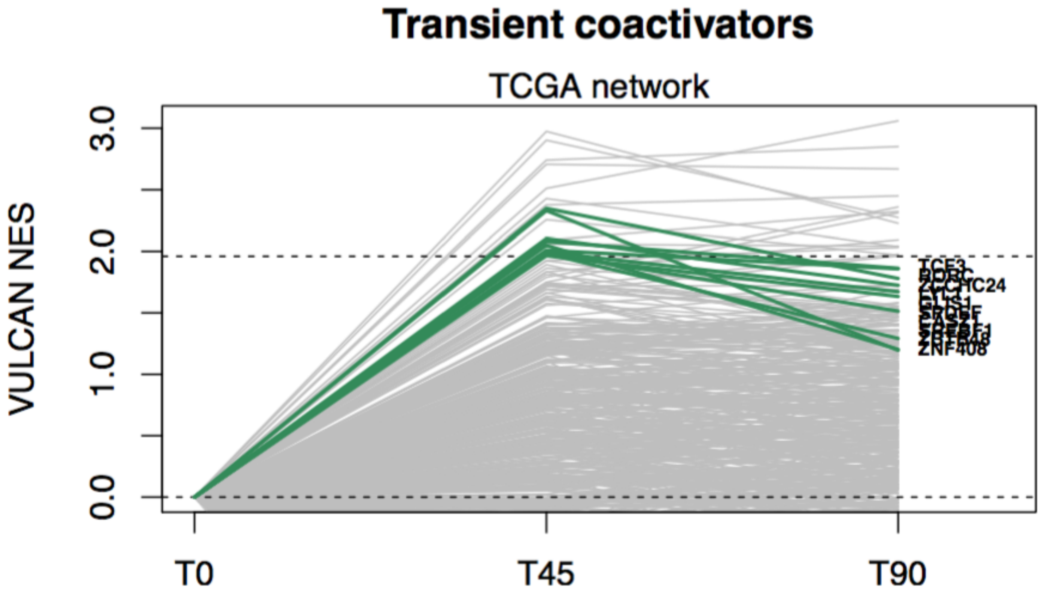


**Fig. S18: Global TF activity after Estradiol Treatment in MCF7 cells, inferred using the TCGA network, highlighting TFs significantly upregulated at 45 minutes but not at 90 minutes.**

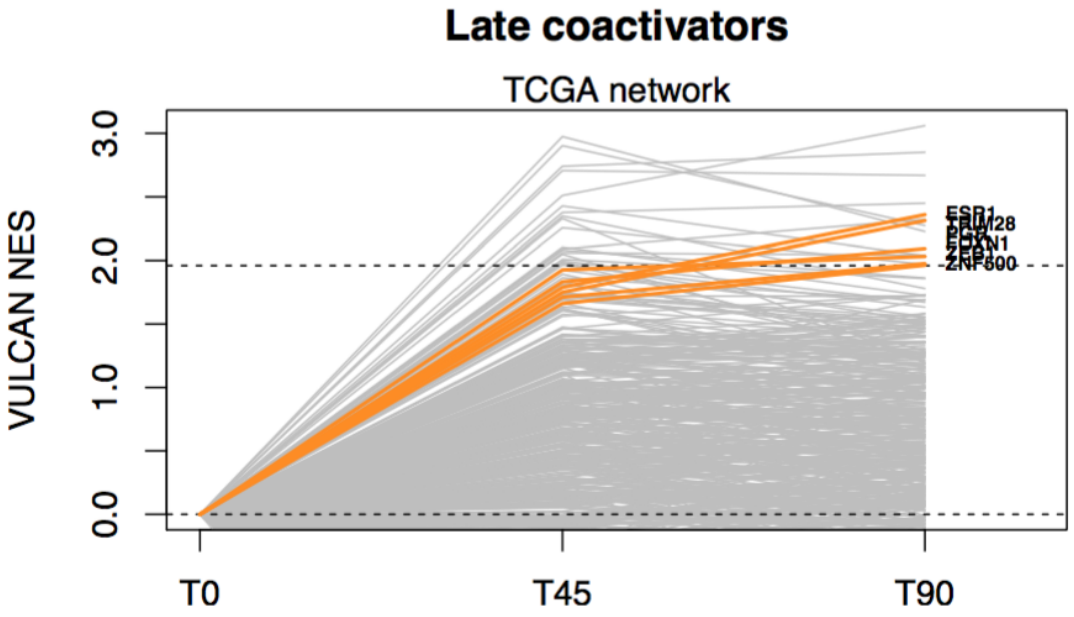


**Fig. S19: Global TF activity after Estradiol Treatment in MCF7 cells, inferred using the TCGA network, highlighting TFs significantly upregulated at 90 minutes but not at 45 minutes.**

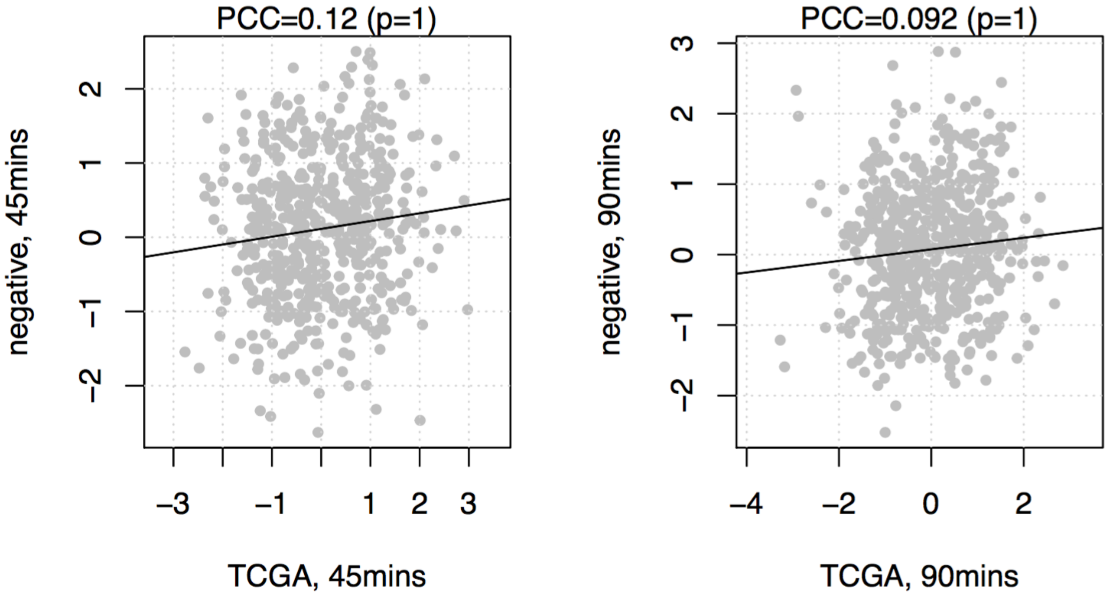


**Fig. S20: Comparison between activities inferred through a breast cancer TCGA dataset and the AML dataset. PCC indicates the Pearson Correlation Coefficient.**

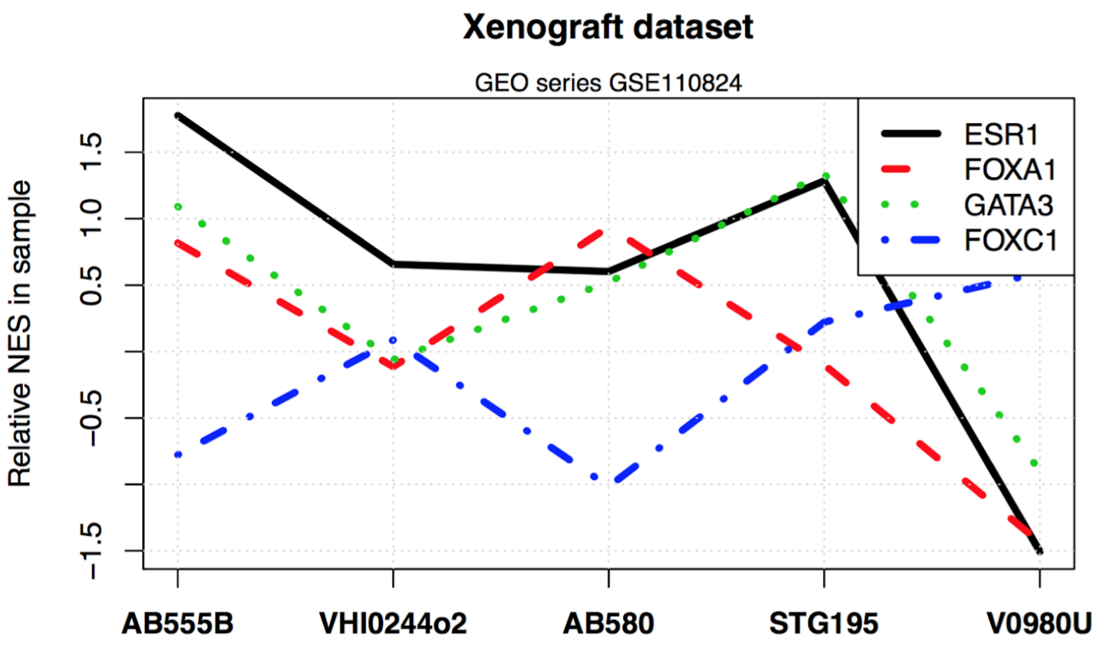


**Fig. S21: VULCAN Activity scores for a few TFs extracted from the ER-targeted ChIP-Seq Xenograft (dataset GSE110824).**

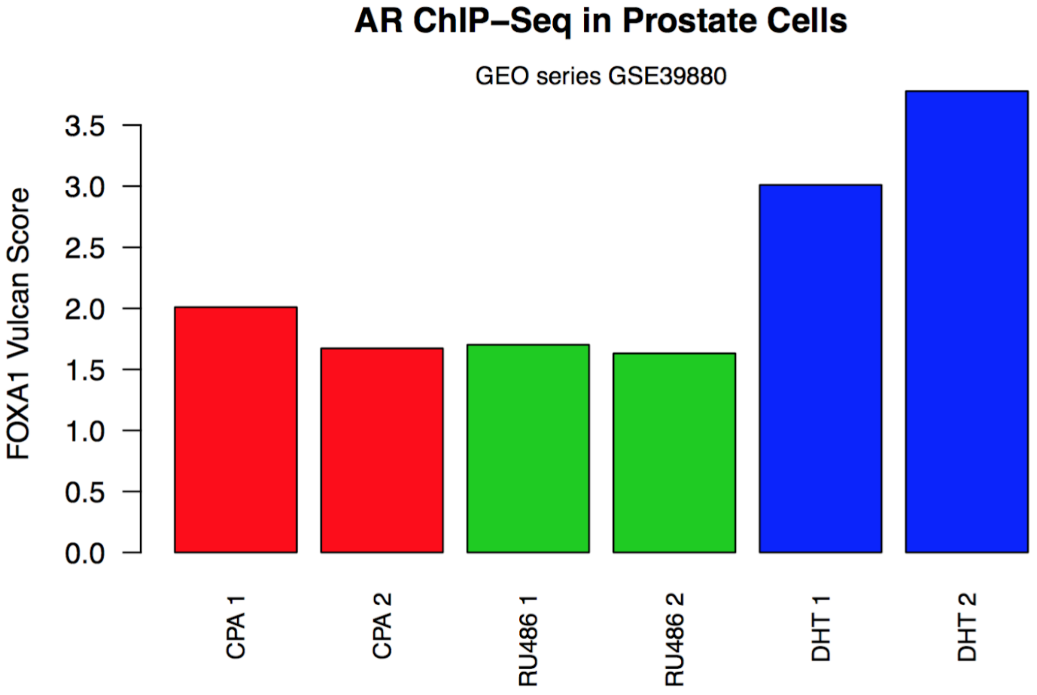


**Fig. S22: VULCAN Activity scores for FOXA1 in Prostate cell lines (dataset GSE39880).**

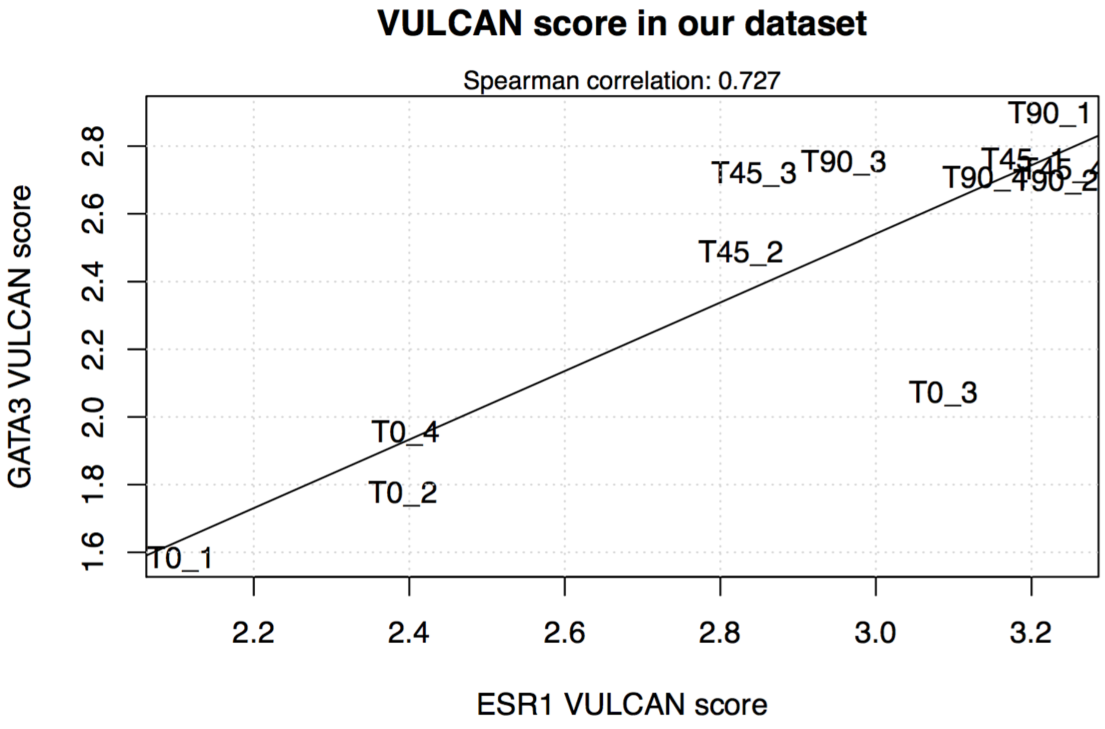


Fig. S23: VULCAN scores of GATA3 and ESR1 in our dataset. Individual samples are indicated.

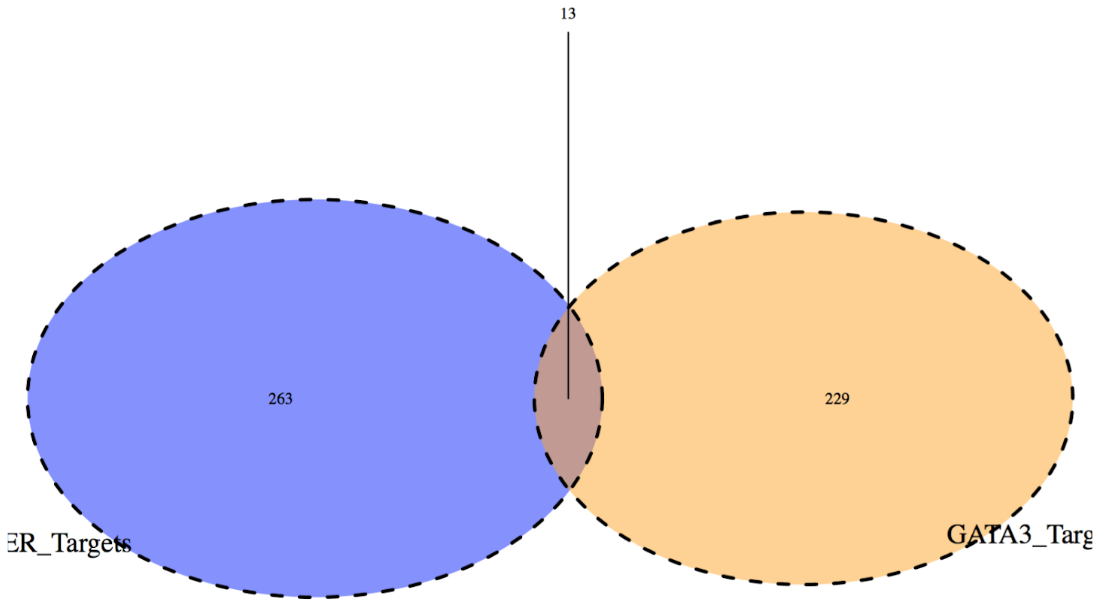


**Fig. S24: Example of target intersection between GATA3 and ER according to the MSigDB database of canonical TF-specific motifs in putative target gene promoters.**

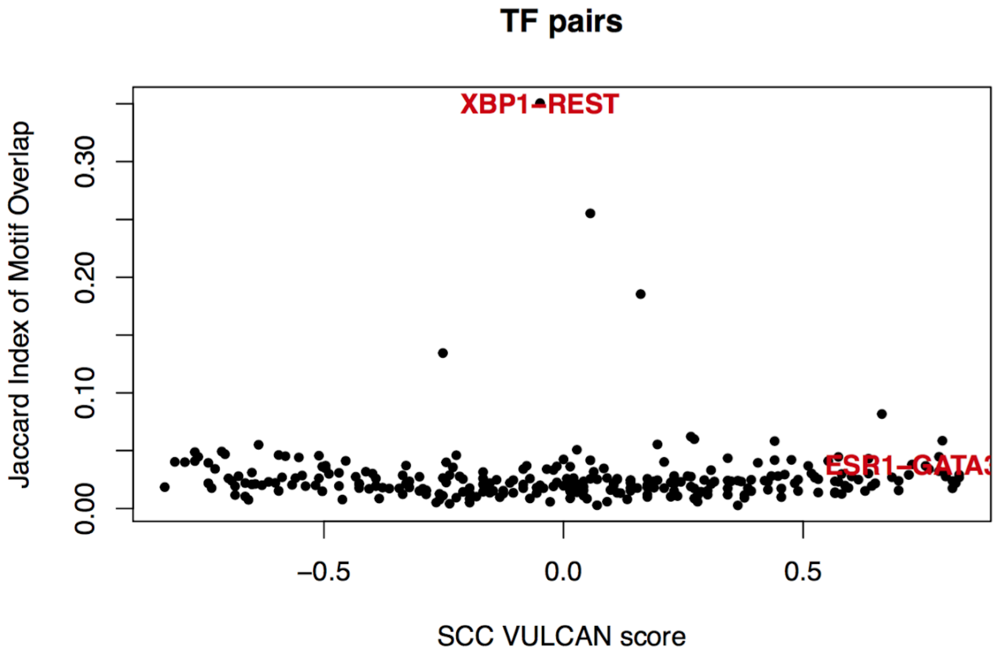


**Fig. S25: TF pairs compared in terms of VULCAN score Spearman Correlation Coefficient in our ER dataset and in terms of Jaccard Index of motif-based target intersection according to the MSigDB C3 collection.**

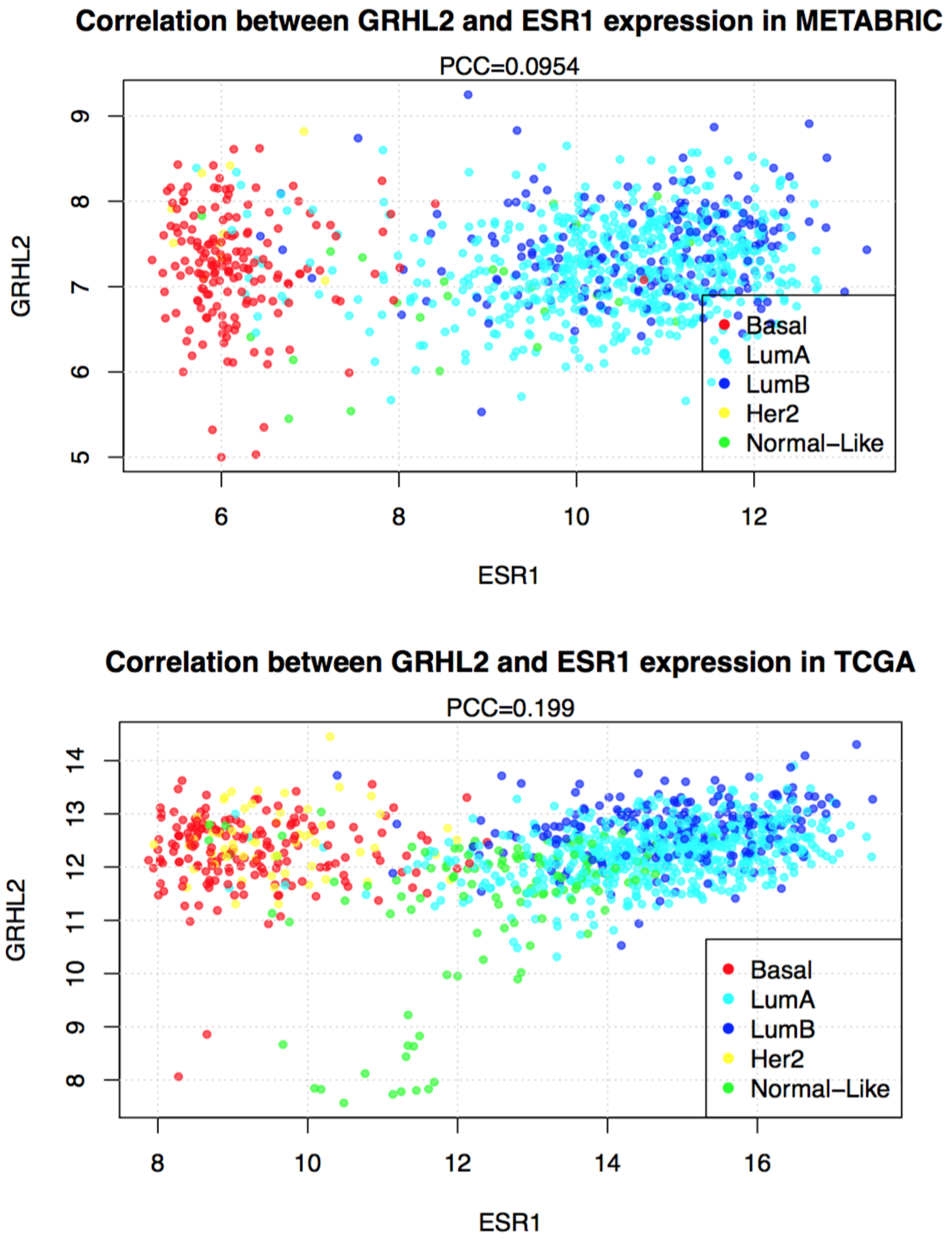


**Fig. S26: Correlation between GRHL2 and ESR1 expression in the TCGA & METABRIC breast cancer datasets.**

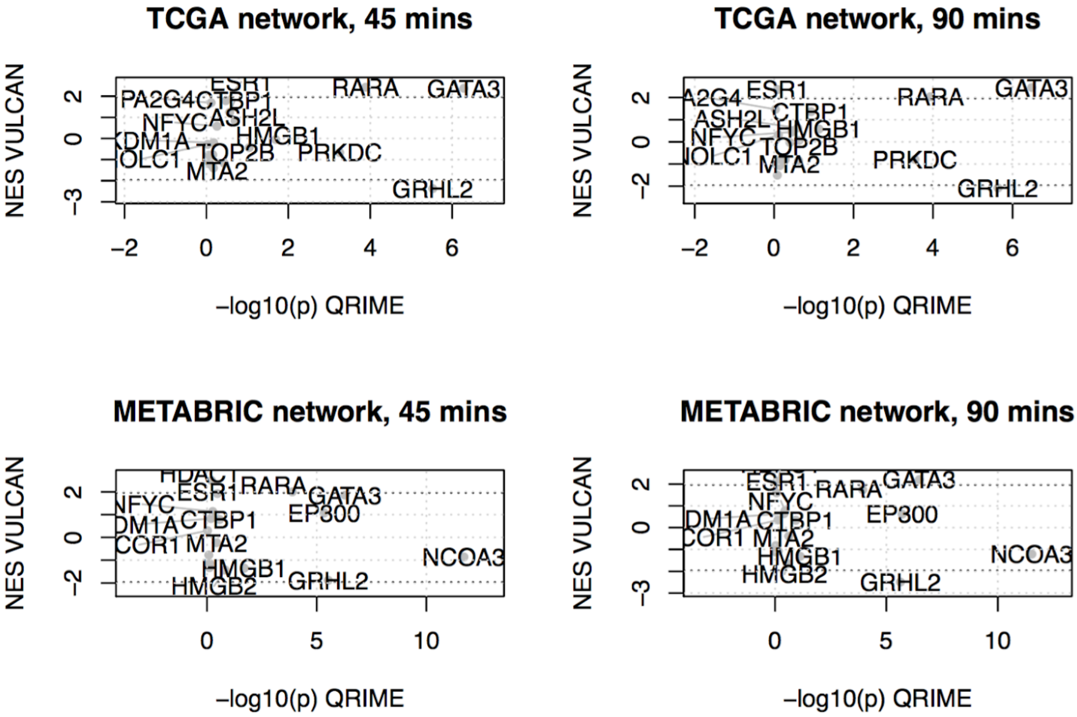


**Fig. S27: Comparison of Normalized Enrichment Score between the QRIME method (x-axis) and the VULCAN method (y-axis) at two time points using two regulatory networks for VULCAN**

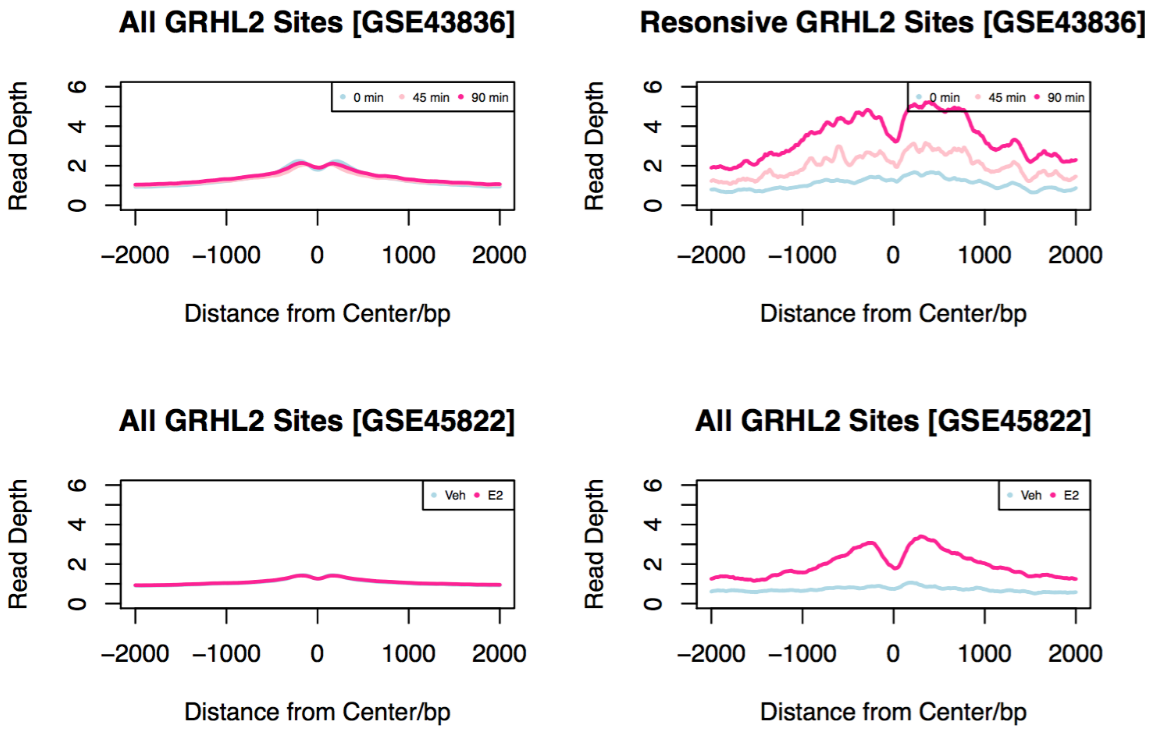


**Fig. S28: Analysis of GRHL2 sites of Gro-Seq data from GSE43836 and GSE45822 both showed that E2 responsive GRHL2 responsive sites are transcriptionally responsive to E2.**

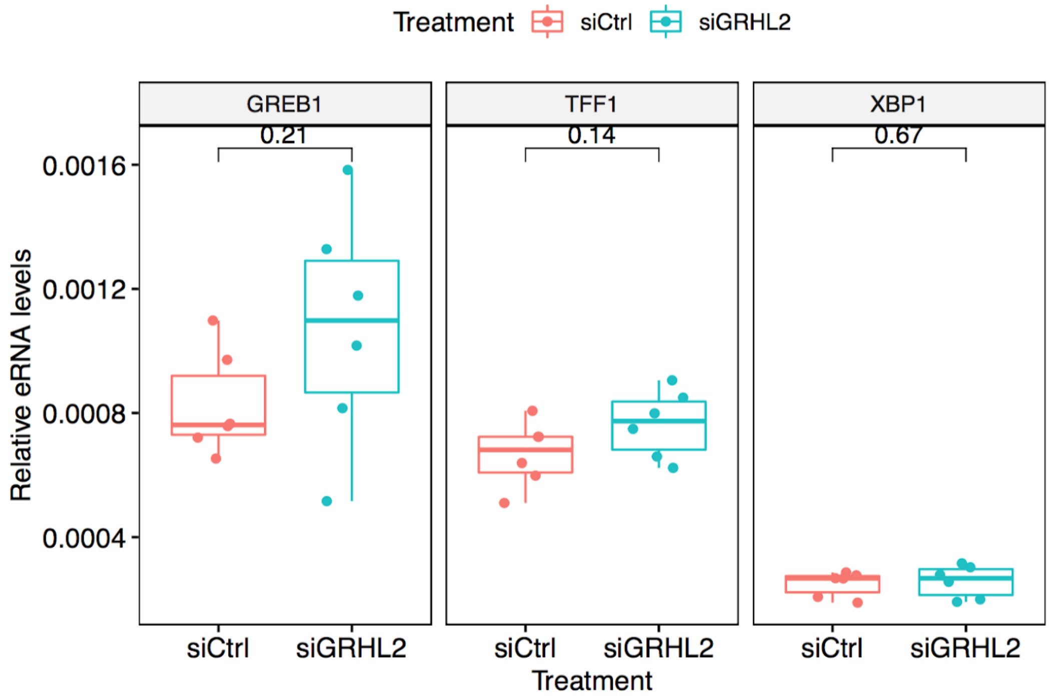


**Fig. S29: Effect of knockdown of GRHL2 on eRNA at E2 responsive binding sites.**

### Supplementary Tables

| | NES, 90' vs 0' | pvalue |
|---|---|---|
| *Bhat esr1 targets not via akt1 up* | 10.3 | 4.6e-25 |
| *Bhat esr1 targets via akt1 up* | 10.3 | 5.3e-25 |
| *Dutertre estradiol response 6hr up* | 8.87 | 7.1e-19 |
| *Frasor response to estradiol up* | 4.93 | 8.3e-07 |
| *Dutertre estradiol response 24hr up* | 4.9 | 9.7e-07 |
| *Stein esr1 targets* | 4.04 | 5.3e-05 |
| *Stein esrra targets responsive to estrogen dn* | 3.74 | 0.00018 |
| *Creighton endocrine therapy resistance 1* | 3.72 | 2e-04 |
| *Stossi response to estradiol* | 3.67 | 0.00024 |
| *Zwang egf interval dn* | 3.55 | 0.00039 |
| *Creighton endocrine therapy resistance 4* | 3.42 | 0.00062 |
| *Pedrioli mir31 targets up* | 3.38 | 0.00071 |
| *Massarweh tamoxifen resistance dn* | 3.24 | 0.0012 |
| *Lein pons markers* | 3.19 | 0.0014 |
| *Reactome hs gag degradation* | 3.19 | 0.0014 |
| *Jiang tip30 targets dn* | 3.17 | 0.0015 |
| *Kegg glycerophospholipid metabolism* | 3.16 | 0.0016 |
| *Geserick tert targets dn* | 3.13 | 0.0018 |
| *Gross hypoxia via hif1a dn* | 3.09 | 0.002 |
| *Valk aml cluster 6* | 3.09 | 0.002 |

**Table S1: aREA results: upregulated MsigDB pathways at 90mins.**

| | NES, 45' vs 0' | pvalue |
|---|---|---|
| *Bhat esr1 targets via akt1 up* | 10.8 | 3e-27 |
| *Bhat esr1 targets not via akt1 up* | 10.4 | 2.8e-25 |
| *Dutertre estradiol response 6hr up* | 8.1 | 5.5e-16 |
| *Frasor response to estradiol up* | 4.54 | 5.6e-06 |
| *Stein esrra targets responsive to estrogen dn* | 4.21 | 2.6e-05 |
| *Dutertre estradiol response 24hr up* | 4.15 | 3.4e-05 |
| *Stein esr1 targets* | 4.03 | 5.6e-05 |
| *Vantveer breast cancer esr1 up* | 3.84 | 0.00012 |
| *Geserick tert targets dn* | 3.69 | 0.00022 |
| *Pedrioli mir31 targets up* | 3.61 | 3e-04 |
| *Reactome glycerophospholipid biosynthesis* | 3.39 | 0.00069 |
| *Zwang egf interval dn* | 3.38 | 0.00072 |
| *Naba ecm affiliated* | 3.31 | 0.00093 |
| *Stossi response to estradiol* | 3.29 | 0.00099 |
| *Lien breast carcinoma metaplastic vs ductal dn* | 3.25 | 0.0011 |
| *Kegg glycerophospholipid metabolism* | 3.25 | 0.0012 |
| *Creighton endocrine therapy resistance 1* | 3.24 | 0.0012 |
| *Reactome hs gag degradation* | 3.21 | 0.0013 |
| *Valk aml cluster 6* | 3.2 | 0.0014 |
| *Massarweh tamoxifen resistance dn* | 3.1 | 0.0019 |

**Table S2: aREA results: upregulated MsigDBpathways at 45mins.**
